# Decoding the multimorbidities among psychiatric disorders and cognitive functioning

**DOI:** 10.1101/837914

**Authors:** E. Golovina, M.H. Vickers, C.D. Erb, J.M. O’Sullivan

**Affiliations:** Liggins Institute, The University of Auckland; A Better Start National Science Challenge; School of Psychology, The University of Auckland

## Abstract

The regulatory contribution that single-nucleotide polymorphisms (SNPs) associated with psychiatric and cognitive phenotypes make to multimorbidity is unknown. Here, we integrate 3D genome organization and expression quantitative trait (eQTL) analyses to identify the genes and biological pathways that are functionally impacted by 2,893 GWAS SNPs associated with cognitive functioning and five psychiatric disorders (i.e. attention deficit hyperactivity disorder (ADHD), anxiety, bipolar disorder (BD), unipolar depression (UD) and schizophrenia (SCZ)). The analysis revealed 33 genes and 62 pathways that were commonly affected by the gene regulatory interactions associated with all six phenotypes despite there being no common SNPs and eQTLs. 38 ADHD-, 78 anxiety-, 81 BD-, 169 UD-, 225 SCZ- and 185 cognition-associated genes represent known drug targets. Four genes were affected by eQTLs from all six phenotypes. Collectively, our results represent the foundation for a shift from a gene-targeted towards a pathway-based approach to the treatment of multimorbid conditions.

## Introduction

Attention-deficit hyperactivity disorder (ADHD), severe anxiety, bipolar disorder (BD), unipolar depression (UD) and schizophrenia (SCZ) are highly prevalent psychiatric disorders^1^. Typically, these disorders are accompanied by cognitive advantages or deficits^2^ that may be associated with increased population level mortality. Epidemiological research has reported that ADHD^3^, anxiety^3, 4^, BD^3^, UD^5^, SCZ^6^ and cognitive functions are multimorbid conditions, suggesting that common biological mechanisms may underlie these phenotypes. Despite evidence for significant genetic heritability^7^, how the underlying genetic architecture of these phenotypes contributes to the observed multimorbidity is incompletely understood. Understanding the regulatory mechanisms underlying these observed multimorbidities will be useful for prognosis, prediction, therapeutic approaches, and understanding drug side-effects. Genome-wide association studies (GWAS) have identified thousands of single nucleotide polymorphisms (SNPs) that are significantly associated with psychiatric disorders and cognitive functions^7^. The majority of these SNPs fall within non-coding genomic regions, consistent with their being expression quantitative trait loci (eQTLs) that are linked to, or enriched within, local or distal regulatory elements^8, 9^. Identifying the functional impact of these SNPs remains a significant hurdle^10^. The three-dimensional (3D) organization of the genome emerges from the sum of the nuclear processes and includes cell type and tissue-specific spatial interactions between regulatory regions and the genes that they control^11^. Recent studies have demonstrated the potential of integrating GWAS SNPs, genome structure and eQTLs for the identification of functional regulatory interactions in SCZ^12, 13^. However, how genetic variants contribute to multimorbidities among psychiatric phenotypes and cognition remains poorly defined.

Here, we integrated data on the 3D genome organization and eQTLs to identify tissue-specific spatial regulatory impacts associated with the development of psychiatric disorders and cognitive functioning (see Methods). The identified regulatory interactions include cis-(<1Mb) and trans-acting (>1Mb) intra- and interchromosomal regulatory connections. eGenes from all phenotypes co-occurred in 62 common biological pathways consistent with the observed multimorbidities among psychiatric and cognitive phenotypes. Drug-eGene interaction analysis identified potential pharmacological influences on these multimorbid phenotypes. Our observations reveal the extent of the shared genetic influences, tissue-specific regulatory mechanisms, biological pathways, and drugs to multimorbidities among ADHD, anxiety, BD, UD, SCZ and cognitive phenotypes. These results highlight novel and existing opportunities for therapeutic drug repurposing to modify psychiatric disorders and cognitive functioning.

## Results

### Functional impacts of genetic variants explain the multimorbidities between psychiatric disorders and cognition

Genetic architecture is a major contributor to the development of psychiatric and cognitive phenotypes^14^. We hypothesized that multimorbidity is driven by genetic variants (e.g. SNPs, structural variants, indels, etc) that regulate tissue-specific expression of genes that co-occur within specific biological pathways and thus affect the phenotype (Fig. 1).

**Fig. 1.**
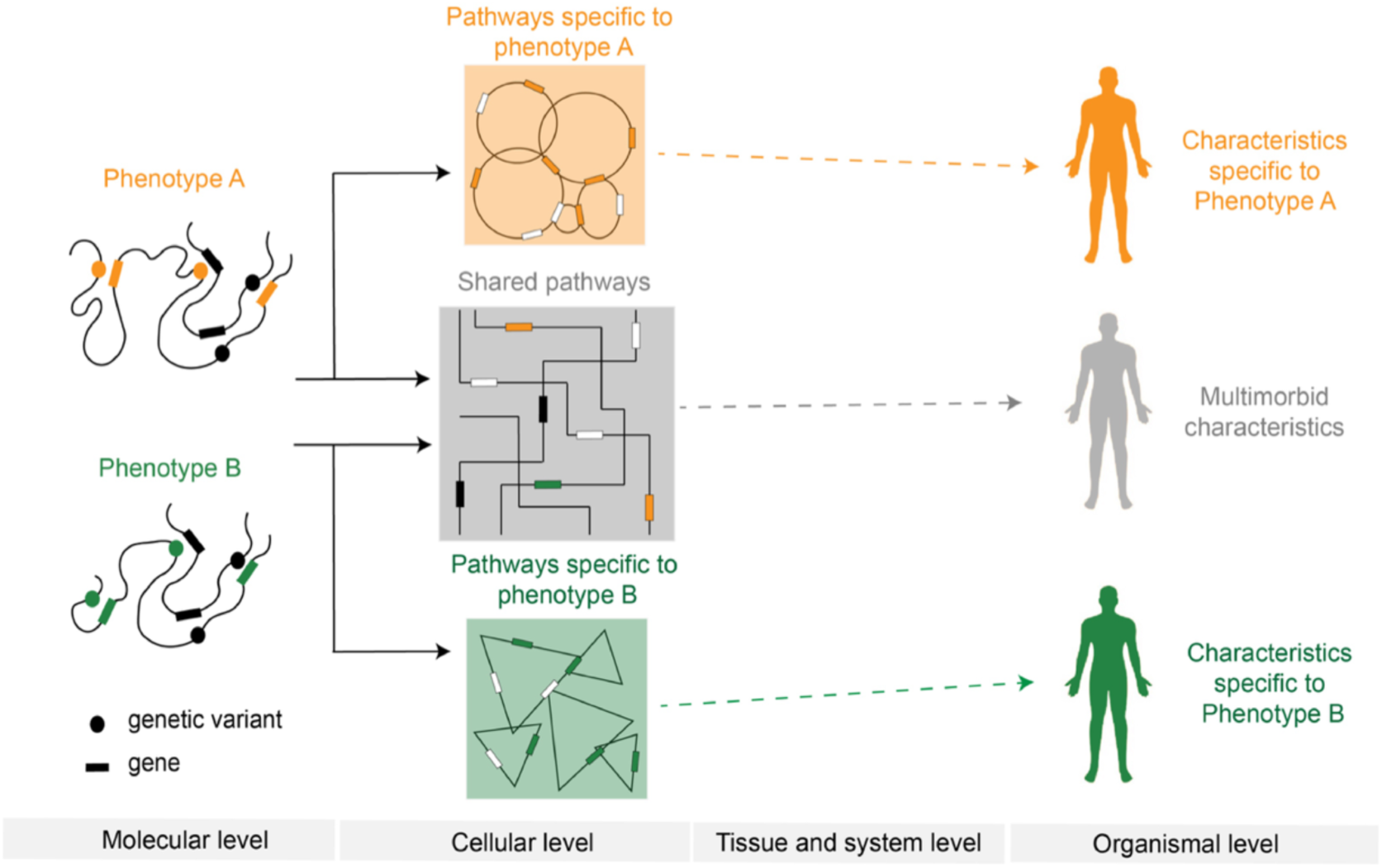
Genetic model of multimorbidity and the SNP-phenotype relationship. Phenotype-specific genetic variants alter tissue-specific gene expression by changing regulatory connections within the 3D dimensional organization of the genome. The gene products, whose expression is altered, interact within biological pathways. Multimorbidity results when affected gene products co-occur within pathways. The co-occurrence of affected gene products within shared pathways changes the way pathways respond to environmental signals and thus affects cellular activities at tissue and system levels.

### SNPs associated with psychiatric disorders and cognition regulate distant genes

SNPs represent the most common type of genetic variation and are widely associated with phenotypes in GWAS^15^. SNPs (n=2,893) associated (p < 1×10^−6^) with cognitive functioning, ADHD, anxiety, BD, UD and SCZ were obtained from the GWAS Catalog (www.ebi.ac.uk/gwas/; Supplementary Table 1). CoDeS3D^16^ was used to integrate GWAS and genome structure data to identify tissue-specific spatial eQTLs for 2,088 (∼70%) of the SNPs (Supplementary Fig. 1 and 2a; GTEx multi-tissue dataset v7, Supplementary Table 3). The 2,088 eQTLs (>40% intronic, >35% intergenic; Supplementary Fig. 2b) were involved in 9,527 statistically significant (FDR < 0.05; Benjamini Hotchberg^17^) SNP-eGene pairs and a total of 45,269 cis- and trans-acting regulatory interactions across 48 different human tissues (Fig. 2, Supplementary Fig. 3, Supplementary Tables 4-10).

**Fig. 2.**
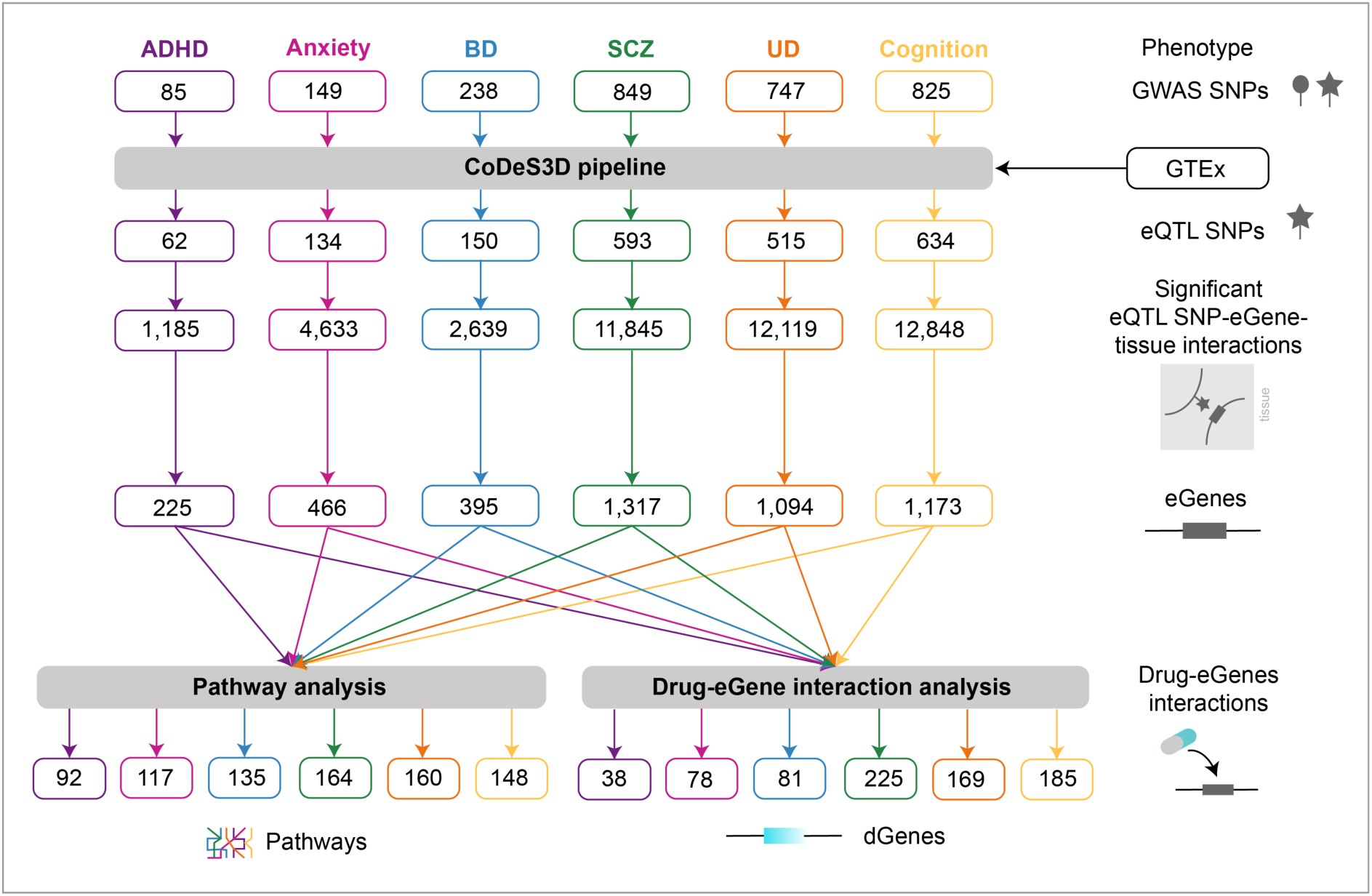
Pipeline used to study the multimorbidities between psychiatric and cognitive phenotypes. SNPs associated with cognition, ADHD, anxiety, BD, SCZ, and UD were obtained from the GWAS Catalog and analysed using CoDeS3D (see Supplementary Fig.1) to identify the genes associated with significant spatial eQTLs. Pathway analysis was used to identify pathways containing co-occurring eGenes for the different phenotypes. Drug-eGene interaction analysis was performed to identify druggable genes (Supplementary Table 4 & 16).

### SNPs and eQTLs are mostly unique to individual phenotypes

We intersected the spatial eQTL sets associated with ADHD, anxiety, BD, UD, SCZ and cognitive functioning to identify shared genetic variation between these phenotypes. There were no SNPs common to all phenotypes (Fig. 3a). Similarly, there were no eQTLs shared by all phenotypes and most were found to be unique to individual phenotypes (Fig. 3b). Among the psychiatric disorders, SCZ and BD show the largest eQTL overlap (69 eQTLs). Only pairwise eQTL overlaps were identified between psychiatric disorders and cognition. No shared eQTLs were identified between ADHD and cognition. Previous studies have also identified SNPs associated with combined phenotypes, i.e. SCZ+cognition^2^, BD+cognition^2^ and BD+SCZ^18^, however, consistent with our findings, most of these phenotype-associated SNPs were individualized. The lack of (or small) SNP and eQTL overlaps among psychiatric and cognitive phenotypes suggests that multimorbidity is less likely due to shared genetic variants, but rather may be explained by their regulatory contributions on genes and pathways.

**Fig. 3.**
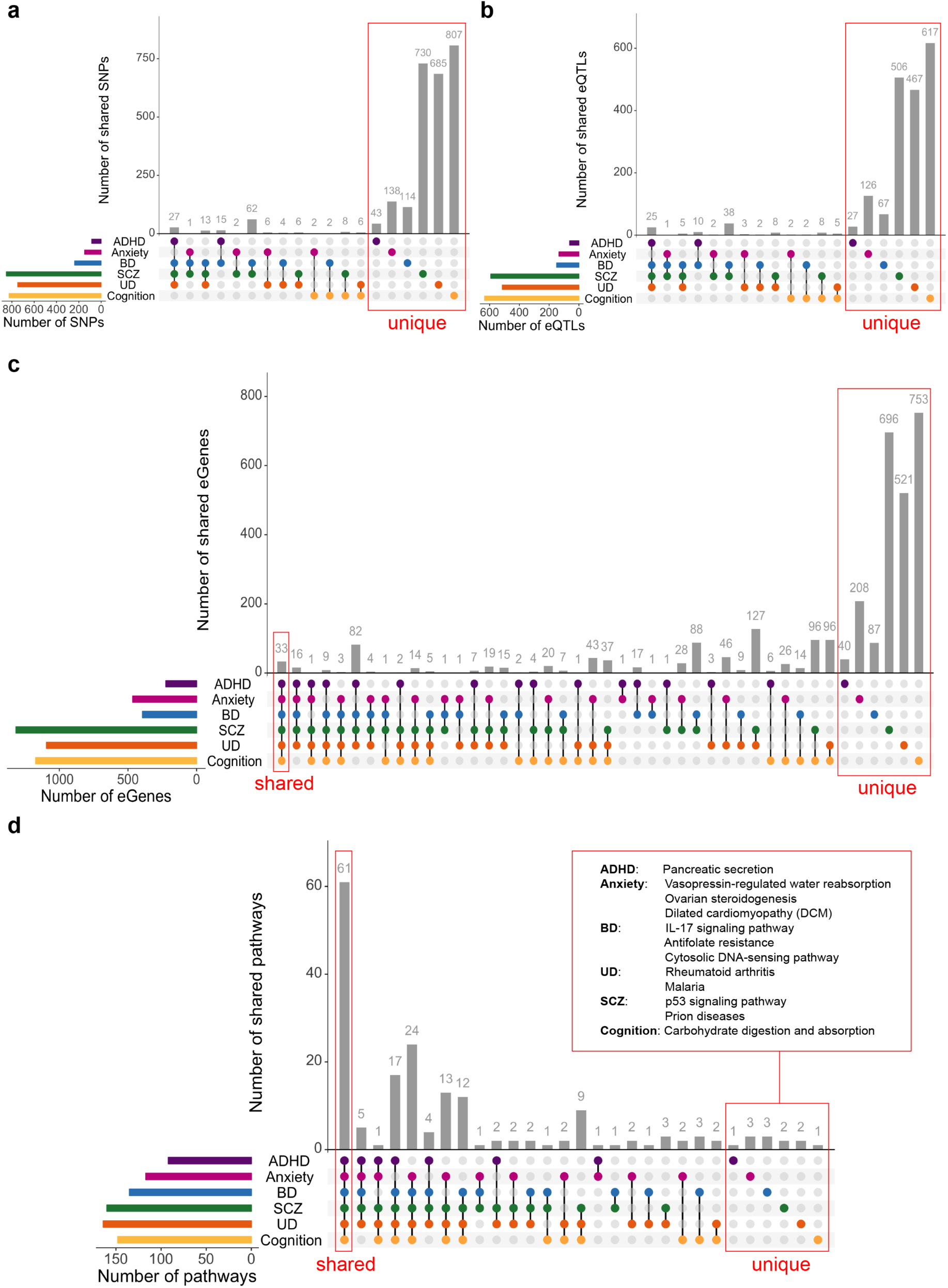
Shared biological pathways link psychiatric disorders and cognition. Psychiatric disorders and cognitive functions have low levels of genetic similarity at the SNPs (**a**), eQTLs (**b**), and eGene (**c**) levels. **d.** Psychiatric disorders and cognition share a large degree of commonality at the biological pathways level. Among the most impacted pathways, 61 were shared between psychiatric disorders and cognition, 66 - across all five psychiatric disorders. Only one pathway (i.e. Pancreatic secretion) was unique to ADHD, 3 pathways (i.e. Vasopressin-regulated water reabsorption, Ovarian steroidogenesis and Dilated cardiomyopathy (DCM)) were specific to anxiety, 3 - to BD (i.e. IL-17 signaling pathway, Antifolate resistance and Cytosolic DNA-sensing pathway), 2 - to SCZ (i.e. p53 signaling pathway and Prion diseases), 2 - to UD (i.e. Rheumatoid arthritis and Malaria) and 1 - to cognition (i.e. Carbohydrate digestion and absorption). The full list of shared and unique pathways is in Supplementary Table 15.

### Psychiatric disorders and cognition share common eGenes

Multimorbidities among psychiatric disorders and cognition could result from the regulatory effects of shared or phenotype-specific eQTLs on common gene targets. Common eGenes (n=33) were identified for eQTLs associated with all six phenotypes (Fig. 3c). Bootstrapping simulations (n=10,000) randomly generating phenotype-associated eGenes confirmed that the observed overlaps were significant (*p* value < 0.01; Supplementary Fig. 4). SCZ and UD shared the greatest number of eGenes (n=374). ADHD and cognition have 58 common eGenes despite having no shared eQTLs (Fig. 3b & 3c).

Expression levels of the 33 common eGenes we identified are associated with cis- and trans-eQTLs from linked variants on chromosomes 3, 6, 10 and 19 (Supplementary Fig. 5, Supplementary Table 11). ADHD, anxiety, BD, SCZ, UD and cognition phenotypes had 1-13 cis-acting eQTLs located within chr3:52256696-53455568 that were associated with the transcript levels of ten eGenes (i.e. *GLYCTK*, *GNL3*, *ITIH4*, *MUSTN1*, *NEK4*, *NT5DC2*, *SFMBT1*, *TMEM110*, *WDR82* and *PBRM1*). Chr6:25177507-32914725 contained a total of 111 eQTLs from the six phenotypes that were associated with the expression of 22 eGenes (including five transcription factors, i.e. *ZFP57*, *ZNF165*, *ZSCAN23*, *ZSCAN31*, *ZSCAN9*). Chr10:103816827-105039240 contained between 3-14 phenotype specific eQTLs associated with transcriptional levels of the *AS3MT* eGene. The majority of phenotype-associated SNPs in these regions are in linkage disequilibrium (D’ > 0.5) (Supplementary Fig. 6 & 7).

Previous studies have identified loci linked to phenotype-related SNPs and genes that they can disrupt^13, 19, 20^. For example, genetic variants at 3p21 region were linked to *GLYCTK*, *GNL3* and *ITIH4* genes that were previously implicated in BD and SCZ^21, 22^. Altered *AS3MT* gene expression (associated with rs7085104 SNP in 10q24.32 locus) in the context of SCZ has recently been reported^13^. The *ITIH3-*rs2535629 and *AS3MT*-rs7085104 associations have also been linked to combined ADHD+BD+UD+SCZ phenotype^20^. However, these assignments are typically based on the assumption that the closest gene to the variant is responsible for the phenotype. Analysis of SNP-eGenes spatial connections showed that only 5-7% are explained by SNP associations with a closest eGene. For example, the intronic *ITIH3-* rs2535629 eQTL SNP (associated with ADHD, BD, UD and SCZ^20^) correlates with expression of the closest (*ITIH3*) gene. However, *ITIH3-*rs2535629 is also involved in spatial regulatory interactions with 12 eGenes (i.e. *ITIH4*, *PPM1M*, *GNL3*, *RAF1, MUSTN1*, *NEK4*, *NT5DC2*, *PBRM1*, *RBMS3*, *SFMBT1*, *TMEM110*, *WDR82*) (Supplementary Table 12). These results suggest that incorporating data on spatial genome organization enables to identify more local and distal eQTL-gene connections that can be missed but may contribute to multimorbid phenotypes.

### Psychiatric disorders and cognition share common biological processes

Gene ontology (GO) analysis (g:Profiler^23^ toolset) of the thirty-three shared eGenes identified significant enrichment (adjusted *p* < 0.05, corrected by the SCS algorithm) in gene expression, transcription, metabolic, biosynthetic and regulatory processes (Supplementary Fig. 8). Notably, ontological analyses of eGenes specific for each phenotype revealed common associations with neurodevelopment (e.g. “nervous system development”), immune system processes (e.g. “immune response”), responses to environmental stimuli and signal transduction (Supplementary Table 13).

### Psychiatric disorders and cognition share a large number of biological pathways

The pleiotropic effects of eQTLs and co-occurrence of the affected eGenes within biological pathways could contribute to the underlying multimorbid conditions (Fig. 1). Pathway analysis using eGenes (shared and specific for each phenotype) identified 61 common biological pathways (Fig. 3d, Supplementary Table 14). These pathways contain a mixture of phenotype shared and specific eGenes (Supplementary Fig. 9) and are associated with human diseases (e.g. alcoholism, glioma, HTLV-I infection), signal transduction (e.g. neurotrophin signaling, Wnt, mTOR, MAPK, cAMP, Ras, thyroid hormone signalling pathways), neurodevelopment (e.g. axon guidance) and learning (e.g. long-term potentiation (LTP)) and immunity (e.g. antigen processing and presentation pathway) (Supplementary Table 15). An additional 66 pathways contain co-occurring eGenes from all of the psychiatric disorders (Fig. 3d, Supplementary Table 15). Bootstrapping analysis (n=10,000) confirmed that these overlaps were significant (*p* value < 0.01; Supplementary Fig. 10).

The neurotrophin signaling pathway, arguably one of the important pathways in developmental neurobiology^24^, contained eGenes associated with eQTLs from all phenotypes (Fig. 4). Most of the eGenes within the neurotrophin signalling cascade were regulated by trans-acting eQTLs (Fig. 4). Dysregulation in the neurotrophin signaling cascade can impact downstream pathways, e.g. axon guidance and LTP, which also contain co-occurring eGenes associated with cognition and psychiatric phenotypes (Supplementary Fig. 11 & 12).

**Fig. 4.**
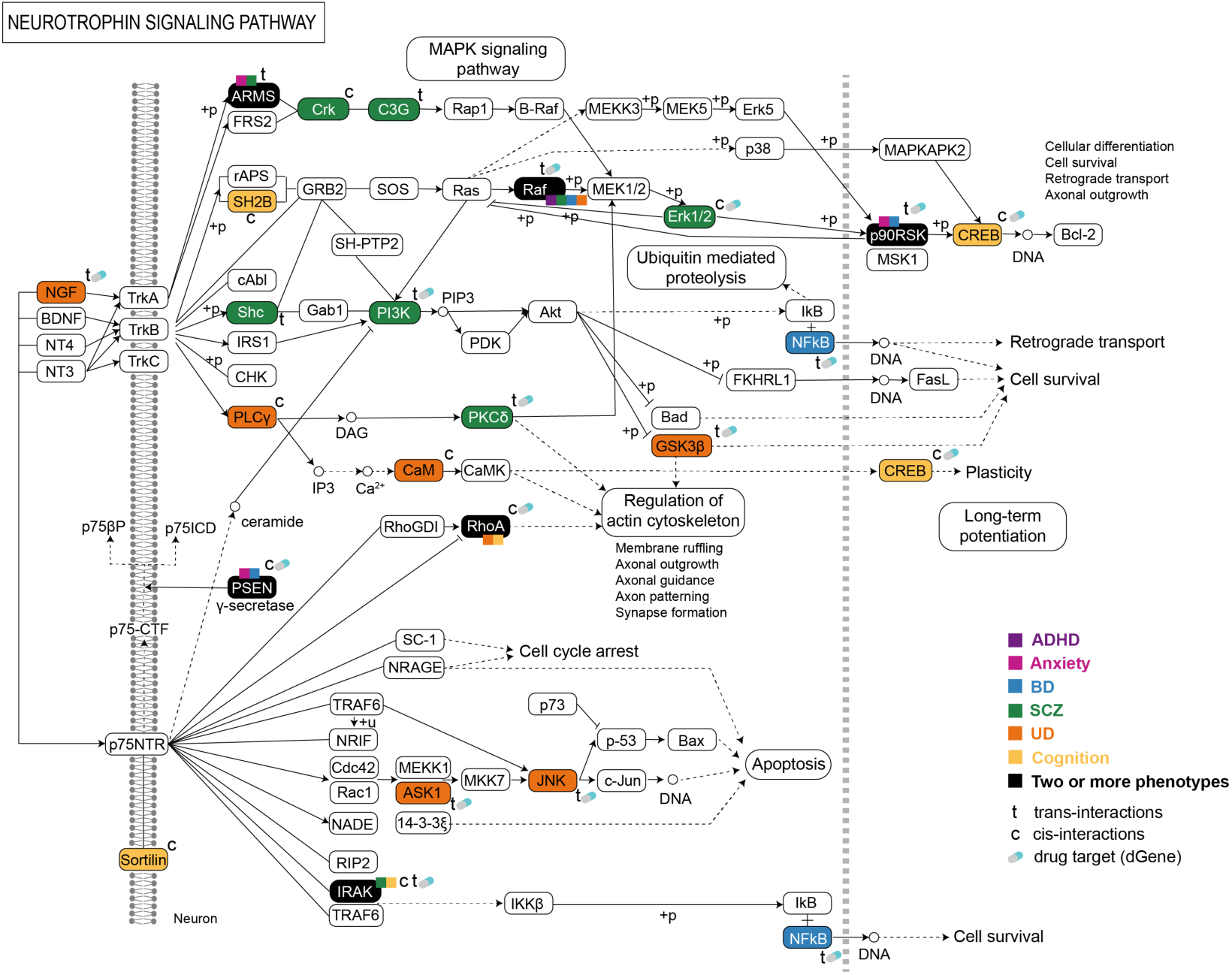
Psychiatric and cognitive SNPs mark eQTLs that associate with gene expression within the neurotrophin signaling pathway. The co-occurrence of the affected shared or phenotype-specific eGenes and imbalance in gene expression within this pathway may lead to a series of cellular functions and events associated with psychiatric and cognitive phenotypes and the multimorbidities between them.

We also identified 12 biological pathways that were impacted by eGenes that were unique to individual psychiatric and cognitive phenotypes. For example, eGenes specific to: 1) ADHD impacted pancreatic secretion; 2) anxiety impacted vasopressin-regulated water reabsorption, ovarian steroidogenesis and dilated cardiomyopathy (DCM); 3) BD impacted IL-17 signalling, antifolate resistance and cytosolic DNA-sensing; 4) SCZ impacted p53 signalling and prion diseases; and 5) UD impacted rheumatoid arthritis and malaria pathways. Carbohydrate digestion and absorption was impacted by eGenes specific to cognition.

### Drug-gene interactions uncover pharmaceutical influences of multimorbidity

Drugs to prevent, stabilize or slow the progress of psychiatric and cognitive conditions often have side effects consistent with known multimorbidities. We queried the Drug Gene Interaction database (DGIdb) to identify eGene products that are targeted by existing drugs (Supplementary Table 16). We identified 16% of the ADHD-risk, 16.7% of the anxiety-risk, 20.5% of the BD-risk, 15.4% of the UD-risk, 17.1% of the SCZ-risk eGenes and 15.8% of the cognition-associated eGenes that represent potential targets for pharmaceutical intervention (Supplementary Table 16).

Four eGenes (i.e. *AS3MT*, *FLOT1*, *HLA-A* and *PBRM1*) that are affected by eQTLs from all tested phenotypes are targets for current drugs (Supplementary Fig. 13). The *AS3MT* product is a target for arsenic compounds, which have neurotoxic effects associated with cognitive dysfunction and mood disorders^25^. Carbamazepine targets *HLA-A* and *FLOT1* products and is used to treat BD, depression and SCZ. Similarly, everolimus targets the *PBRM1* product, which is currently being explored as a therapeutic option for neurological diseases characterized by cognitive impairments^26^. Everolimus improves cognitive function while aggravating depression and anxiety^26^.

Notably, the products of these shared druggable eGenes are involved in the same overlapping biological pathways. For example, *FLOT1* products are enriched in insulin signaling while *PBRM1* products are involved in hepatocellular carcinoma pathway. The products of *HLA-A* are enriched in pathways related to antigen processing and presentation, natural killer cell mediated cytotoxicity, HTLV-I and human papillomavirus infections, cellular senescence. Collectively, these results provide insights into potential drug side-effects associated with multimorbidities among psychiatric disorders and cognition.

### Spatial regulatory eQTL effects are tissue-specific

Brain structural changes are commonly considered as relevant factors in the development of psychiatric and cognitive phenotypes. However, these phenotypes are associated with physiological changes at the level of the whole body, e.g. with impaired functioning of endocrine^27^, immune^28, 29^ and cardiometabolic^30^ systems. Therefore, tissue-specific spatial regulatory interactions of eQTL SNPs with their target genes may be important in understanding the multimorbidities among psychiatric and cognitive phenotypes. There was widespread distribution of cis- and trans-acting eQTL regulatory interactions we identified across 48 GTEx tissues (Supplementary Fig. 14). The number of eQTL SNP-eGene interactions acting over distances of <1Mb and ≥1Mb correlated with tissue sample size (Supplementary Fig. 15). By contrast, the numbers of interchromosomal interacting eQTLs in each tissue did not correlate with tissue sample size (Supplementary Fig. 15). Notably, we detected more <1Mb and ≥1Mb-acting eQTL-eGene interactions in thyroid and brain cerebellum, than predicted (Supplementary Fig. 15) for the phenotypes we tested (Supplementary Table 18). This finding is consistent with the previously reported role of the thyroid in the development of cognitive functions and psychiatric disorders^27, 31^.

## Discussion

Psychiatric and cognitive phenotypes were found to share relatively little common genetic risk at the level of SNPs and eQTLs. However, our observations are consistent with the known multimorbidities being explained by the combinatorial action of eGenes within shared biological pathways. We identified 57 distinct patterns of multimorbidity between psychiatric disorders and cognition at the pathways level with 61 biological pathways being shared across all six phenotypes. Only 12 biological pathways that were impacted by eGenes were unique to specific phenotypes. Collectively, our results suggest that emerging interactions between eQTLs, genes and pathways at the tissue and system levels may be potential drivers of multimorbidity among psychiatric disorders and cognition.

Increasing evidence indicates that infections during pregnancy, at birth and in early childhood increase the risk of ADHD^32^, BD^33^ and SCZ^33^. We identified shared pathways associated with infectious diseases (e.g. HTLV-I infection, hepatitis B) as containing eGenes regulated by shared and condition specific eQTLs. Notably, we identified the immune candidate genes *IRAK1* and *IRAK3*^34^ as being associated with SCZ eQTLs. Collectively, our observations are consistent with psychiatric phenotypes having inflammatory components^35, 36^ to their development albeit the exact mechanisms remain unknown.

Studies on psychiatric disorders suggest that they are multisystem conditions^37–39^. This can be rationalized by dysfunctions in other organs signalling for changes in the brain. This is epitomized by hypo and hyperthyroidism, where altered thyroid hormone supplies cause changes in brain functioning, leading to psychiatric disorders (e.g. SCZ, BD, anxiety and depression)^27, 40^. Interestingly, we found common pathways associated with thyroid hormone functioning (e.g. thyroid hormone signalling pathway) across all six phenotypes. Dysregulation and imbalanced gene expression within this pathway may be associated with impaired neurodevelopment^40^.

It remains unclear when the observed multimorbidities arise during development. Are they cause or consequence of the psychiatric and cognitive conditions with which they associate? The genes and pathways that we identified are consistent with a strong developmental component to these phenotypes. For example, axon guidance, Wnt, GnRH, Hippo, TGF-beta, thyroid hormone and neurotrophin signaling pathways were affected by eQTLs specific for each phenotype and these pathways are central to neurodevelopment. Similarly, neurotrophins have been implicated in neuroplasticity, learning and memory through receptor signalling. Therefore, the identification of a cascade of eQTLs associated with the expression of genes affected in psychiatric (i.e. *CRK*, *RAPGEF1*, *PIK3R1*, *PRKCD*, *SHC4*, *MAPK3*) and cognitive (i.e *SORT1*, *SH2B1* and *ATF4*) phenotypes within the neurotrophin signalling pathway is consistent with a central role in the development of these multimorbid conditions. Further analysis of the shared pathways and tissue-specific drug interactions with potential targetable gene products will provide insights into potential opportunities for therapeutic intervention. Additionally, here we focused on gene products that could be pharmacologically modulated, but the use of technologies such as gene therapy will expand the list of potential interventions.

Psychiatric disorders and cognitive functions are complex multifactorial phenotypes whose development depends on complex, often non-linear, dynamic interplay between genetic, epigenetic and environmental factors. Given this, our study has several limitations. Firstly, we’re focusing on the GWAS tag-SNPs only, without capturing the underlying causal SNPs within a ‘tagged’ LD block, which can limit the functional inferences from GWAS^10, 41^. Secondly, increasing the number of GWAS studies on certain phenotypes (e.g. ADHD, anxiety, etc.) will result in the identification of additional novel SNP loci. Thirdly, SNPs do not explain all of the estimated heritability in psychiatric and cognitive phenotypes, suggesting that other factors (e.g. rare variants, indels, structural variation, methylation, etc.) also contribute^14^. Fourthly, the GTEx eQTL data used in this study was largely from European individuals aged 40 years and older^42^. Thus, robust predictions of the regulatory mechanisms of psychopathology and cognitive functioning in different populations and developmental stages requires additional data sets. Also, the cognitive functions we investigated here were diverse (with 791 out of 825 SNPs being associated with intelligence, see Supplementary Table 1) and further research is needed to look more precisely at specific aspects of cognitive functioning. Finally, the GTEx tissue and Hi-C datasets were not paired. This confounds the study of tissue-specific regulatory interactions particularly when combined with potential cellular senescence effects during the GTEx resampling procedure. Despite this, our analysis provides a starting point for further mechanistic and functional investigation. Replication analyses will increase the robustness of the results and provide a clearer indication of cross-phenotype overlap and phenotype-specific genetic architecture.

In conclusion, we have described new pathways for multimorbidity and identified drug interactions that may be clinically relevant for the treatment and prevention of psychiatric and cognitive phenotypes as well as enhancing and facilitating health outcomes. Collectively, our analyses inform the extent of shared genetic influences, tissue-specific regulatory mechanisms, and pathways between complex multi-genic phenotypes. Our results provide the basis for a change away from a gene centric approach to therapy, instead identifying pathways for the treatment multimorbid psychiatric and cognitive conditions. Future applications of the spatial genetic approach we used will cross the molecular, cellular, tissue and system levels to define personalized multimorbid disease risk profiles, therapeutic targets and drug side-effects. This approach can be extended systematically to other multimorbid conditions.

## Online Methods

### GWAS SNPs

Single-nucleotide polymorphisms (SNPs) associated with five psychiatric disorders (i.e. ADHD, anxiety, BD, UD and SCZ) were downloaded from the NHGRI-EBI GWAS Catalog (www.ebi.ac.uk/gwas/; 07/12/2018) with *p* values <1×10^-6^ (see Data and code availability). SNPs associated with nine cognitive functions (i.e. intelligence, information processing speed, cognition, reading ability, reasoning, mathematical ability, infant expressive language ability, language performance and speech perception) were downloaded from the NHGRI-EBI GWAS Catalog (www.ebi.ac.uk/gwas/; 14/07/2018) with *p* values <1×10^-6^ (see Data and code availability). Phenotypes for SNPs were defined as the traits associated with SNPs in the NHGRI-EBI GWAS Catalog. Functional annotation of SNPs was performed using wANNOVAR tool^43, 44^. Genomic positions and annotations of SNPs were obtained for the human genome build hg19 release 75 (GRCh37) (see Data and code availability).

### Hi-C data and data processing pipeline

In total, 28 Hi-C chromatin interaction libraries were used in this study (see Supplementary Table 4). We used high-resolution Hi-C chromatin interaction libraries from seven cell lines (GM12878, HMEC, HUVEC, IMR90, K562, KBM7 and NHEK)^45^. Hi-C interaction data were downloaded from GEO (see Supplementary Table 4) using the download_default_data module of the CoDeS3D pipeline. Additionally, we requested and downloaded Hi-C raw data for neurons, other tissues and cell lines from GEO and dbGaP databases (see Supplementary Table 4). We received the access approval for HeLa (project #18446: “Finessing predictors of cognitive development (part 2)”) and cortical plate and germinal zone neurons (project #16489: “Finessing predictors of cognitive development”) Hi-C data from the dbGaP database. We analysed the raw data as outlined in Rao et al. (Juicer^46^, version 1.5) to generate Hi-C libraries. The pipeline included BWA alignment of paired-end reads onto the hg19 reference genome, merging paired-end read alignments and removing duplicates. The resulting and previously prepared files containing cleaned Hi-C contacts locations (i.e. *_merged_nodups.txt files) were further processed to get Hi-C chromatin interactions libraries in the following format: read name, str1, chr1, pos1, frag1 mapq1, str2, chr2, pos2, frag2, mapq2 (str = strand, chr = chromosome, pos = position, frag = restriction site fragment, mapq = mapping quality score, 1 and 2 correspond to read ends in a pair). Reads where both ends had a mapq ≥ 30 were included in the final library.

### CoDeS3D pipeline

The CoDeS3D^16^ pipeline was used to identify genes that spatially interact with phenotype-associated eQTL SNPs (Supplementary Fig. 1). To identify DNA fragments, the hg19 reference genome was digested with the same restriction enzyme as used in Hi-C library preparation (i.e. Mbol or HindIII) using the digest_genome module of CoDeS3D. The process_inputs module was used to identify SNPs location within the DNA fragments. The find_interactions module identified restriction fragments that interact with SNP-containing fragments in each of 28 Hi-C chromatin interaction libraries. The find_genes module identified spatial SNP-gene pairs where SNP-containing fragments interact with fragments overlapping with a gene region. GENCODE transcript model version 19 was used as the reference for gene annotations. The find_eqtls module queried the GTEx database (https://gtexportal.org/, GTEx multi-tissue dataset v7, Supplementary Table 4), with the SNP-gene pairs, to identify cis- and trans-acting eQTL SNP-eGees interactions (i.e. genes, whose tissue-specific expression changes are associated with a SNP). Lastly, in the produce_summary module, the Benjamini-Hochberg FDR control algorithm^17^ was applied to adjust the *p* values of eQTL associations and identify significant eQTL SNP-eGene-tissue interactions (FDR < 0.05).

### Bootstrapping analysis

To estimate that the observed spatial eGenes associations and overlaps are not random we performed bootstrap test with *N*=10000 iterations. On each iteration step, bootstrap samples containing random eGenes were generated for each phenotype (based on the number of eGenes associated with each phenotype). For each bootstrapped overlap the number of shared eGenes was calculated. We defined an overlap as the number of shared eGenes we observe between two or more specific phenotypes (e.g. eGene overlap between ADHD and Anxiety, or eGene overlap among ADHD, Anxiety, BD, UD, SCZ and Cognition). After 10,000 iterations we counted those iterations where the number of shared eGenes in the bootstrapped overlap (*eGenes_bootstrapped*) is greater than or equal to the number of shared eGenes in the observed overlap (*eGenes_observed*). The *p* value was calculated as the sum of these iterations divided by the total number of iterations *N*,

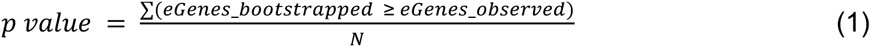

Statistical significance level was determined as 0.01. In other words, if for eGene overlap we estimate *p* value < 0.01, we reject the null hypothesis (our assumption that the observed eGene overlap is due to chance) and accept the alternative hypothesis that the observed relationship is true and not random.

The bootstrap test with *N*=10000 iterations was also performed to estimate that the observed pathway overlaps are non-random. The same procedure was used as for eGenes bootstrapping. Total number of KEGG pathways is 536 (iPathwaysGuide, 09/13/2019).

### LD analysis

LD analysis between the psychiatric disorders and cognition-associated eQTL SNPs from three putative regulatory regions was performed using LDlink 3.7^47^. The LDmatrix module of the tool was used to calculate linkage disequilibrium statistics for eQTL SNPs in all population groups (GRCh37/hg19 genome assembly; SNP RS numbers based on dbSNP151; genotyping data from phase 3 (version 5) of the 1000 Genome Project; African, Ad Mixed American, East Asian, European and South Asian populations.)

### Functional enrichment analysis

Functional gene enrichment analysis was performed using g:GOSt core tool of the g:Profiler^23^ tool set. Three Gene Ontology terms (i.e. biological process, molecular function and cellular component) were used to test if specific categories or biological functions were enriched among the identified eGenes of the regulatory eQTL SNPs. All known human genes were chosen as a statistical domain scope. The significance of the overrepresented GO terms was corrected by the SCS algorithm^23^ (adjusted *p* < 0.05).

### Pathway analysis

Lists of eGenes regulated by phenotype-associated eQTL SNPs were analyzed using Advaita Bio’s iPathwayGuide (https://www.advaitabio.com/ipathwayguide, 09/13/2019) to identify the most impacted biological pathways enriched in psychiatric disorders and cognition. The iPathwaysGuide considers the role, position and relationships of each gene within a pathway that significantly reduces the number of false positives and results in identification of the truly impacted pathways. The FDR algorithm^17^ was applied to correct *p* values for multiple testing and determine significance at the pathway level (FDR < 0.05).

### Correlation analysis

To measure the statistical association between GTEx tissue sample size and the number of cis- and trans-acting eQTL-eGenes interactions we performed Pearson’s correlation analysis (Data and code availability).

### Drug-eGene interaction analysis

We queried the Drug Gene Interaction database^48^ (DGIdb) via the DGIdb API to get information on drugs, their effects and mechanisms of action on the target eGene products.

### URLs

Juicer 1.5: https://github.com/aidenlab/juicer

NHGRI-EBI GWAS Catalog: https://www.ebi.ac.uk/gwas/

wANNOVAR: http://wannovar.wglab.org/

GEO database: https://www.ncbi.nlm.nih.gov/geo/

dbGaP database: https://www.ncbi.nlm.nih.gov/gap/

CoDeS3D pipeline: https://github.com/Genome3d/codes3d-v1

GTEx Portal: https://gtexportal.org/home/

LDlink 3.725: https://ldlink.nci.nih.gov/

g:Profiler (version e95_eg42_p13_f6e58b9): https://biit.cs.ut.ee/gprofiler/

Advaita Bio’s iPathwayGuide (09/13/2019): https://www.advaitabio.com/ipathwayguide

DGIdb (v3.0.2 - sha1 ec916b2): http://www.dgidb.org/

## Data availability

The GWAS Catalog associations (version 1.0.1) were downloaded from https://www.ebi.ac.uk/gwas/ and are available in Supplementary Table 1. The Hi-C datasets that support the findings are available from GEO and dbGaP databases. Accession numbers of these datasets are given in Supplementary Table 2. The GTEx dataset v7 (dbGaP accession phs000424.v7.p2) was used in this study (see Supplementary Table 3). Human genome build hg19 release 75 (GRCh37) was downloaded from ftp://ftp.ensembl.org/pub/release-75/fasta/homo_sapiens/dna/Homo_sapiens.GRCh37.75.dna.primary_assembly.fa.gz. SNP genomic positions (CoDeS3D SNP database) were obtained from ftp://ftp.ncbi.nih.gov/snp/organisms/human_9606_b151_GRCh37p13/. Gene annotations were downloaded from https://storage.googleapis.com/gtex_analysis_v7/reference/gencode.v19.transcripts.patched_contigs.gtf. The datasets generated by the CoDeS3D pipeline that support the findings of this study are available in Supplementary Tables 5-10 and at https://github.com/Genome3d/psychiatric_and_cognitive_multimorbidities/results/codes3d_output. Source data underlying Figs. 2, 3a-d and Supplementary Figs. 2a-b, 3-10, 13-15 are also available at https://github.com/Genome3d/psychiatric_and_cognitive_multimorbidities/data. Supplementary Tables 1-18 are available in figshare with the identifier https://doi.org/10.17608/k6.auckland.10282580.v1.

## Code availability

CoDeS3D pipeline is available at https://github.com/Genome3d/codes3d-v1. All python and R scripts used for data analysis and visualisation are available at https://github.com/Genome3d/psychiatric_and_cognitive_multimorbidities. R version 3.5.2 and RStudio version 1.1.463 were used for all R scripts. All python scripts are based on Python 2.7.15

## Acknowledgements

The authors would like to thank the Genomics and Systems Biology Group (Liggins Institute) for useful discussions. EG is the recipient of a Liggins Ph.D. scholarship. This work was funded by a University of Auckland FRDF grant (Confirming spatial connections to unravel how SNPs affect phenotype; 3714499) to JOS. JOS is also funded by a MBIE Catalyst grant (The New Zealand-Australia LifeCourse Collaboration on Genes, Environment, Nutrition and Obesity (GENO); UOAX1611); Royal Society of New Zealand Marsden Fund [Grant 16-UOO-072]. The Genotype-Tissue Expression (GTEx) Project was supported by the Common Fund of the Office of the Director of the National Institutes of Health, and by NCI, NHGRI, NHLBI, NIDA, NIMH, and NINDS.

The authors wish to acknowledge the use of New Zealand eScience Infrastructure (NeSI) high performance computing facilities, consulting support and/or training services as part of this research. New Zealand’s national facilities are provided by NeSI and funded jointly by NeSI’s collaborator institutions and through the Ministry of Business, Innovation & Employment’s Research Infrastructure programme. URL https://www.nesi.org.nz.

## Author contributions

EG performed the analyses and wrote the manuscript. MV co-supervised EG and commented on the manuscript. CE aided results interpretation and commented on the manuscript. JO supervised EG and co-wrote the manuscript.

## Competing Interests statement

The authors declare no competing interests.

## Supplementary Tables

**Supplementary Table 1:** All GWAS SNPs used in this study

**Supplementary Table 2:** The Hi-C datasets were downloaded from GEO or dbGaP databases and their accession numbers

**Supplementary Table 3:** eQTL associations were based on GTEx multi-tissue dataset v7 (dbGaP Accession phs000424.v7.p2)

**Supplementary Table 4:** The summary of the findings from CoDeS3D, pathway and drug-eGene interaction analyses

**Supplementary Table 5:** Detailed information on spatial eQTL SNP-eGene-tissue interactions associated with ADHD

**Supplementary Table 6:** Detailed information on spatial eQTL SNP-eGene-tissue interactions associated with anxiety

**Supplementary Table 7:** Detailed information on spatial eQTL SNP-eGene-tissue interactions associated with BD

**Supplementary Table 8:** Detailed information on spatial eQTL SNP-eGene-tissue interactions associated with UD

**Supplementary Table 9:** Detailed information on spatial eQTL SNP-eGene-tissue interactions associated with SCZ

**Supplementary Table 10:** Detailed information on spatial eQTL SNP-eGene-tissue interactions associated with cognitive functions

**Supplementary Table 11:** eGenes shared across all psychiatric disorders and cognition

**Supplementary Table 12:** rs2535629 SNP-gene associations linked to psychiatric disorders and cognitive functions

**Supplementary Table 13:** Gene Ontology enrichment analysis

**Supplementary Table 14:** Pathways associated with psychiatric disorders and cognitive functions

**Supplementary Table 15:** Unique and shared pathways between psychiatric disorders and cognitive functions (57 multimorbidity patterns (i.e. phenotypic combinations) and 6 unique patterns).

**Supplementary Table 16:** DGIdb drug-eGene interactions associated with psychiatric disorders and cognitive functions.

**Supplementary Table 17:** dGenes in pathways associated with psychiatric disorders and cognitive functions

**Supplementary Table 18:** Number of cis-(< 1Mb) and trans-acting (≥ 1Mb and interchromosomal) eQTL SNP-eGene interactions per GTEx tissue.

## Supplementary Figures

**Supplementary Fig. 1.**
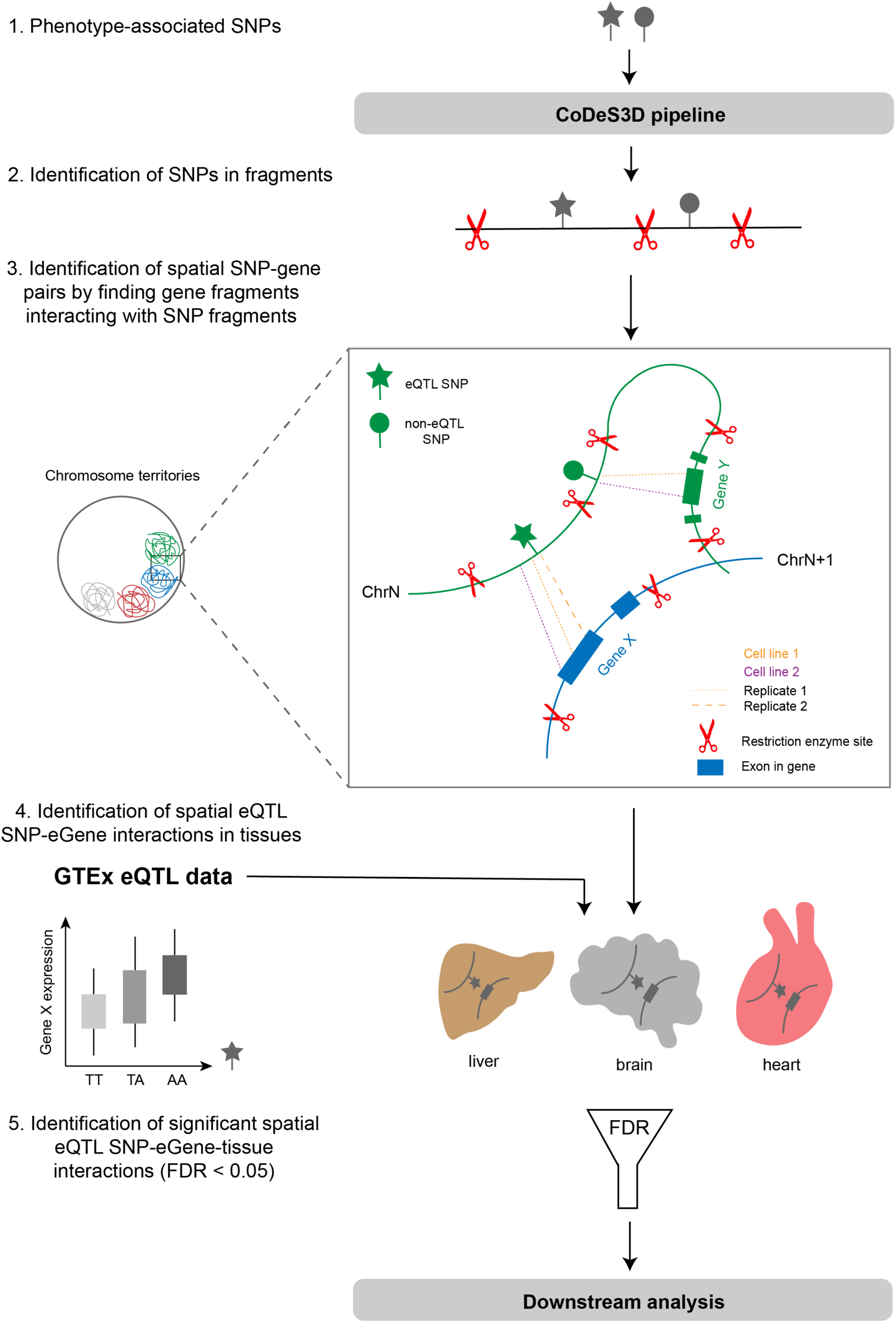
The CoDeS3D pipeline used in this study. Restriction fragments containing SNPs associated with psychiatric disorders and cognition were identified. Hi-C libraries were interrogated to identify genes in fragments that spatially interact (in cis- and trans-) with SNP fragments. The resulting spatial SNP-gene pairs were used to query GTEx database to determine functional tissue eQTL interactions between SNP and eGene (i.e. gene, whose expression is in eQTL with a SNP). Only statistically significant (FDR < 0.05) eQTL SNP-eGene-tissue interactions were further used in the downstream analysis (i.e. functional gene enrichment, pathway analysis, and drug-eGene interactions).

**Supplementary Fig. 2.**
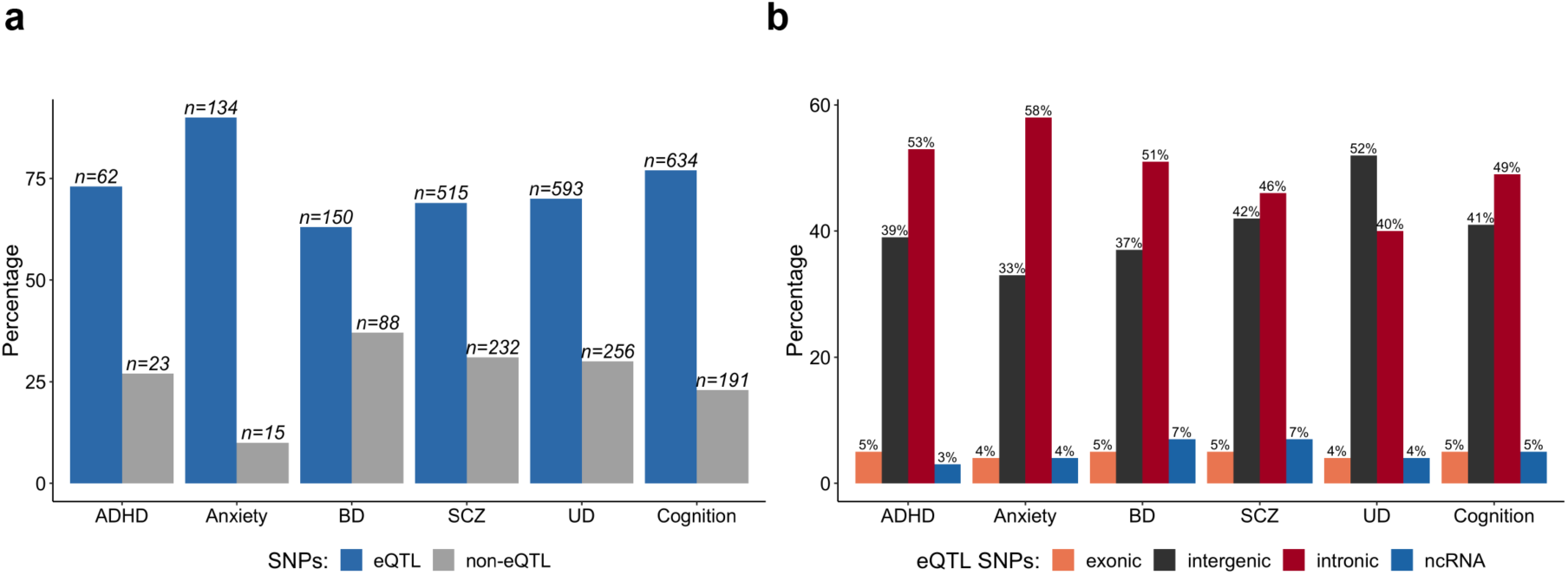
The percentage contribution of GWAS SNPs that are correlated with mRNA expression of genes (i.e. eQTL SNPs) and their distribution in the genome. **a** The majority of the GWAS SNPs associated with psychiatric disorders and cognition impact on gene expression as eQTLs. **b** Most of these eQTL SNPs fall within non-coding regions (i.e. introns and intergenic regions).

**Supplementary Fig. 3.**
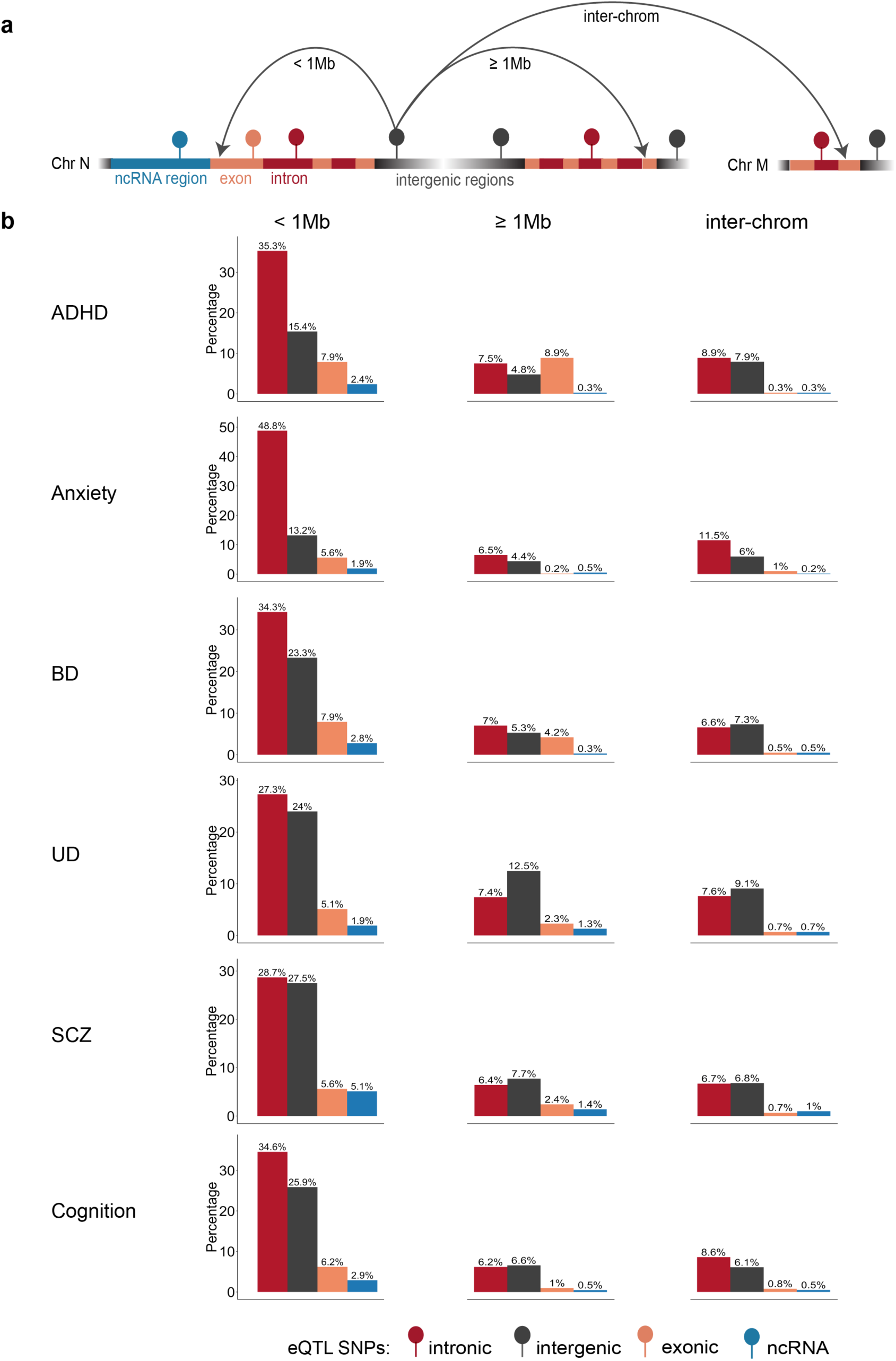
Functional types of eQTL SNPs associated with psychiatric disorders and cognition and the percentage distribution of these eQTL SNP types in the genome. a Schematic representation of cis-(< 1Mb) and trans-acting (≥ 1Mb and interchromosomal) SNPs. b Most of eQTL SNPs were enriched in intronic and intergenic regions and mark regions that regulate expression of genes in close proximity (i.e. in cis-manner). Functional annotation of trans-acting interactions showed that trans-eQTLs involve more coding SNPs (8.9%) in ADHD compared to other phenotypes. Trans-acting regulation in SCZ and UD is associated more with intergenic eQTLs than with intronic regulatory variants. The majority of interchromosomal eQTL effects are intronic and intergenic across all phenotypes

**Supplementary Fig. 4.**
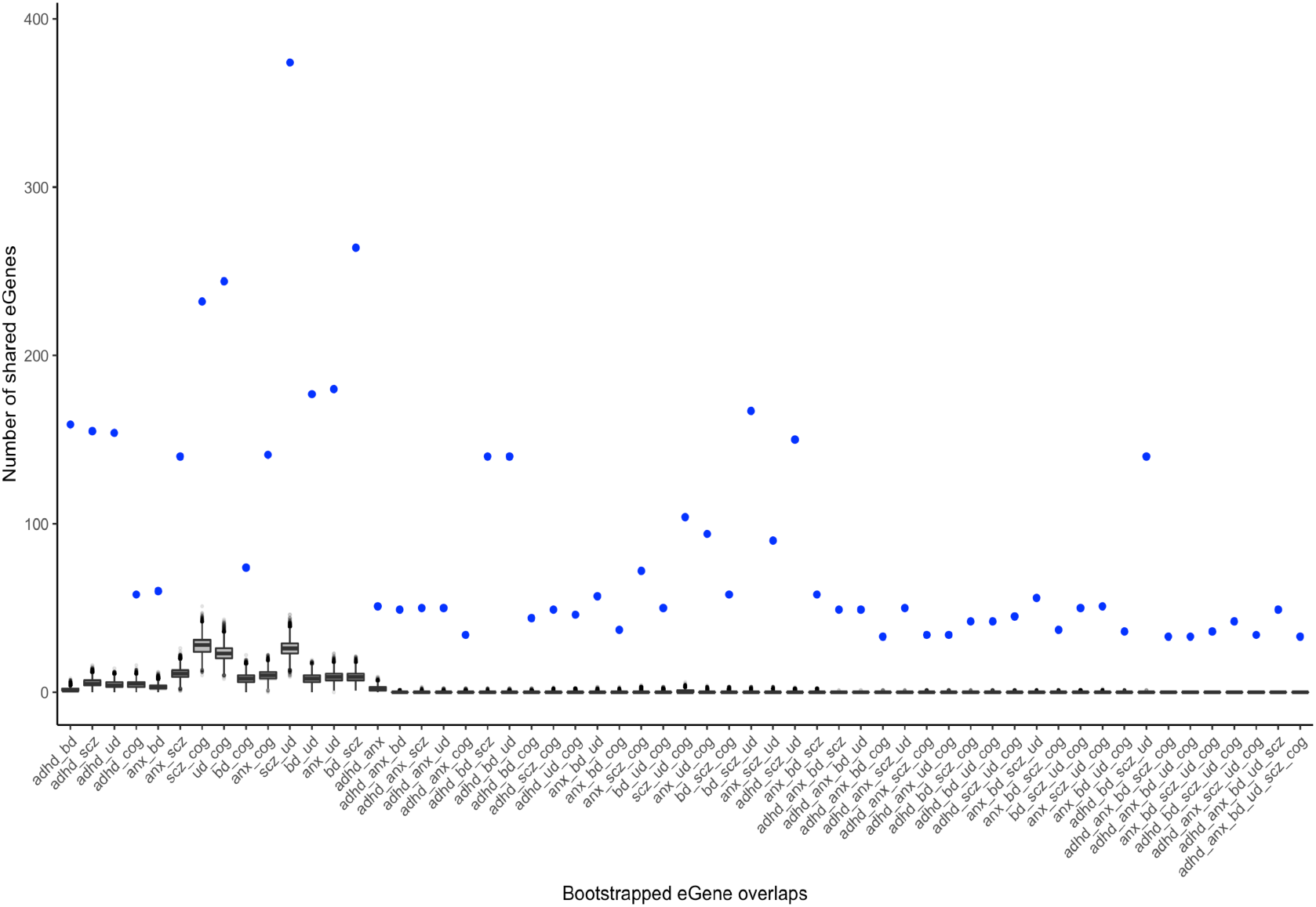
Box plots of the bootstrapped (10,000 iterations) eGene overlaps among psychiatric disorders and cognition. Blue dots indicate the number of shared eGenes in the observed overlaps. The bootstrap test shows that the observed eGenes overlaps are statistically significant (*p* < 0.01) and didn’t occur due to chance.

**Supplementary Fig. 5.**
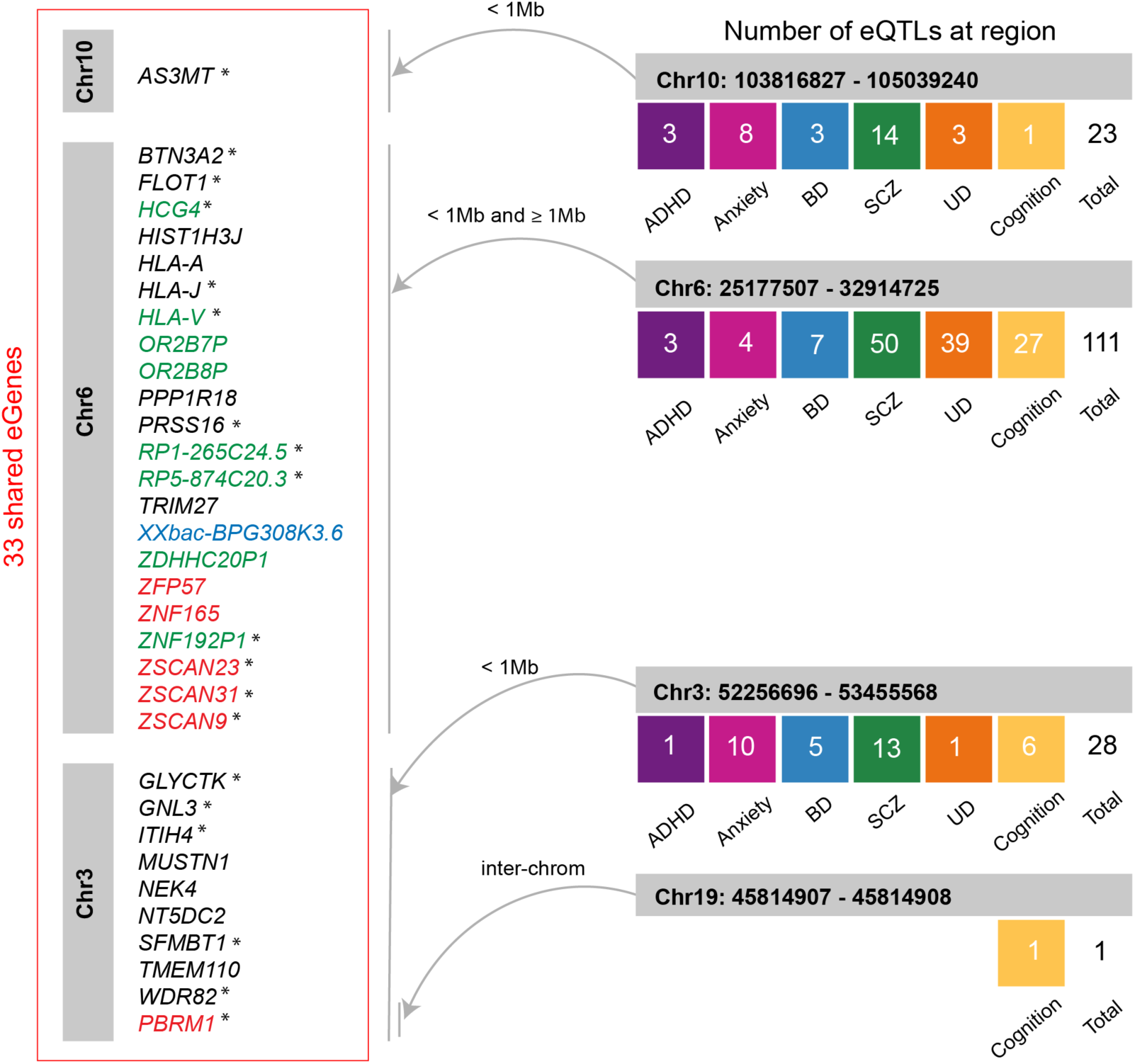
Common eGenes are affected by different eQTLs. We identified thirty-three shared eGenes between psychiatric disorders and cognition. These are located on chromosomes 3, 6 and 10 and regulated in cis- and trans-by multiple eQTL SNPs from four putative regions on chromosomes 3, 6, 9 and 10. Most of the shared eGenes are protein-coding (colored in black). Six of them encode transcription factors (colored in red). eGenes colored in green indicate pseudogenes. One eGene (colored in blue) encodes ncRNA. eGenes marked with asterisks are associated with up- or downregulation in the brain.

**Supplementary Fig. 6.**
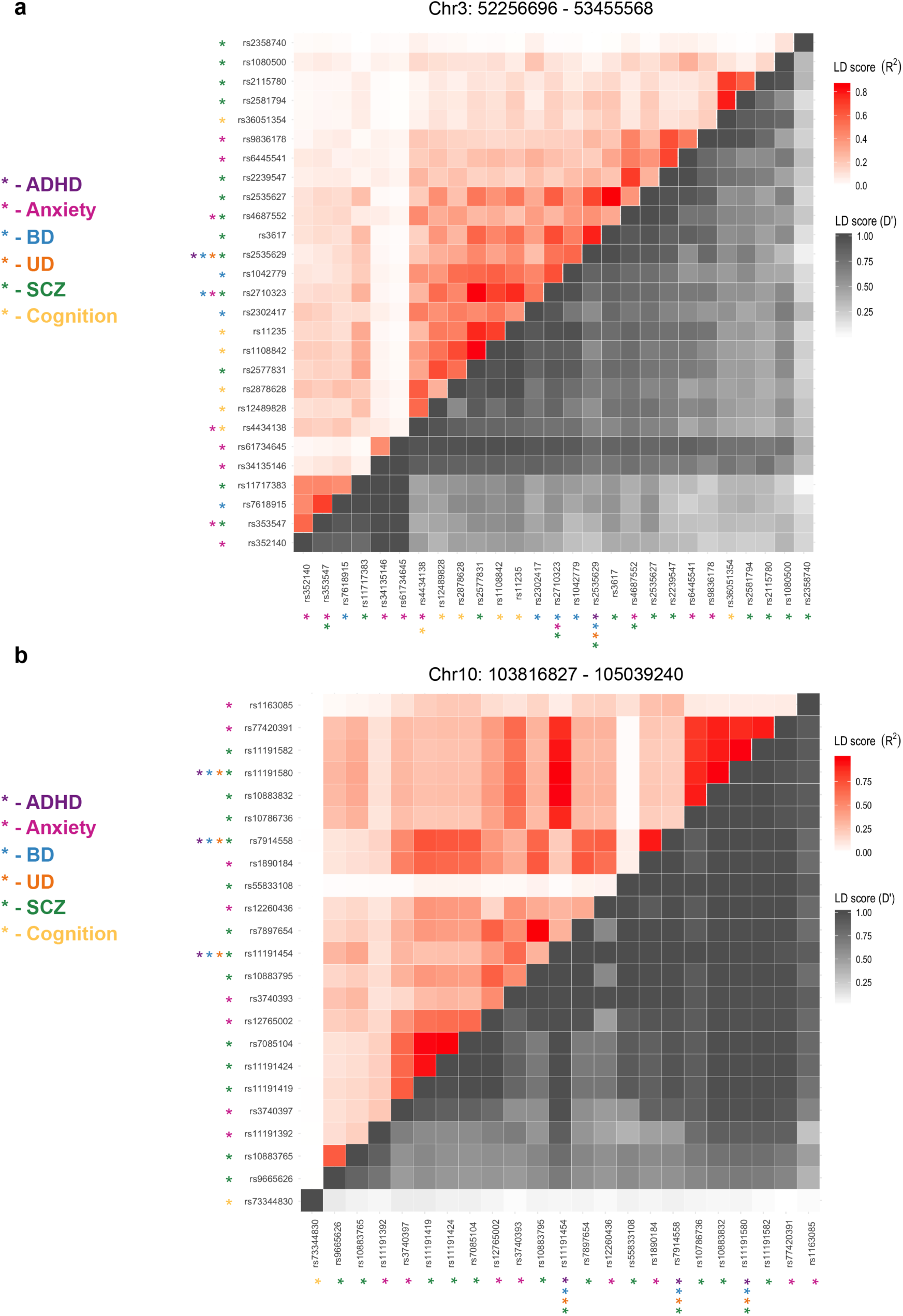
LD plots (R^2^ and D’) of eQTLs associated with psychiatric disorders and cognition and located in putative regulatory regions on chromosome 3 (**a**) and chromosome 10 (**b**). The colors represent the strength of pairwise LD according to R^2^ (red) and D’ (grey) metrics. The colored asterisks mark the SNPs associated with the corresponding phenotypes.

**Supplementary Fig. 7.**
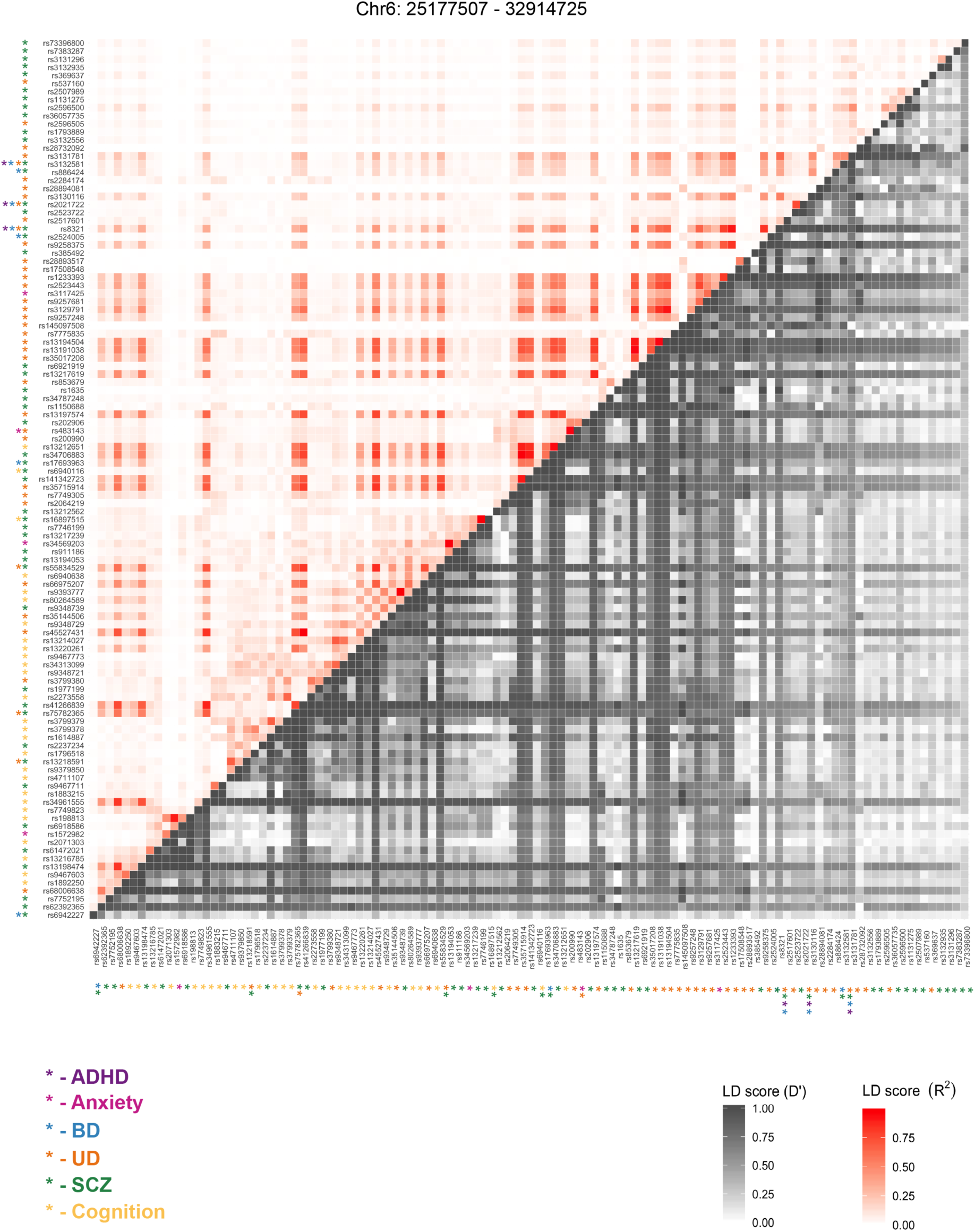
LD plot (R^2^ and D’) of eQTLs associated with psychiatric disorders and cognition and located in putative regulatory regions on chromosome 6. The colors represent the strength of pairwise LD according to R^2^ (red) and D’ (grey) metrics. The colored asterisks mark the SNPs associated with the corresponding phenotypes.

**Supplementary Fig. 8.**
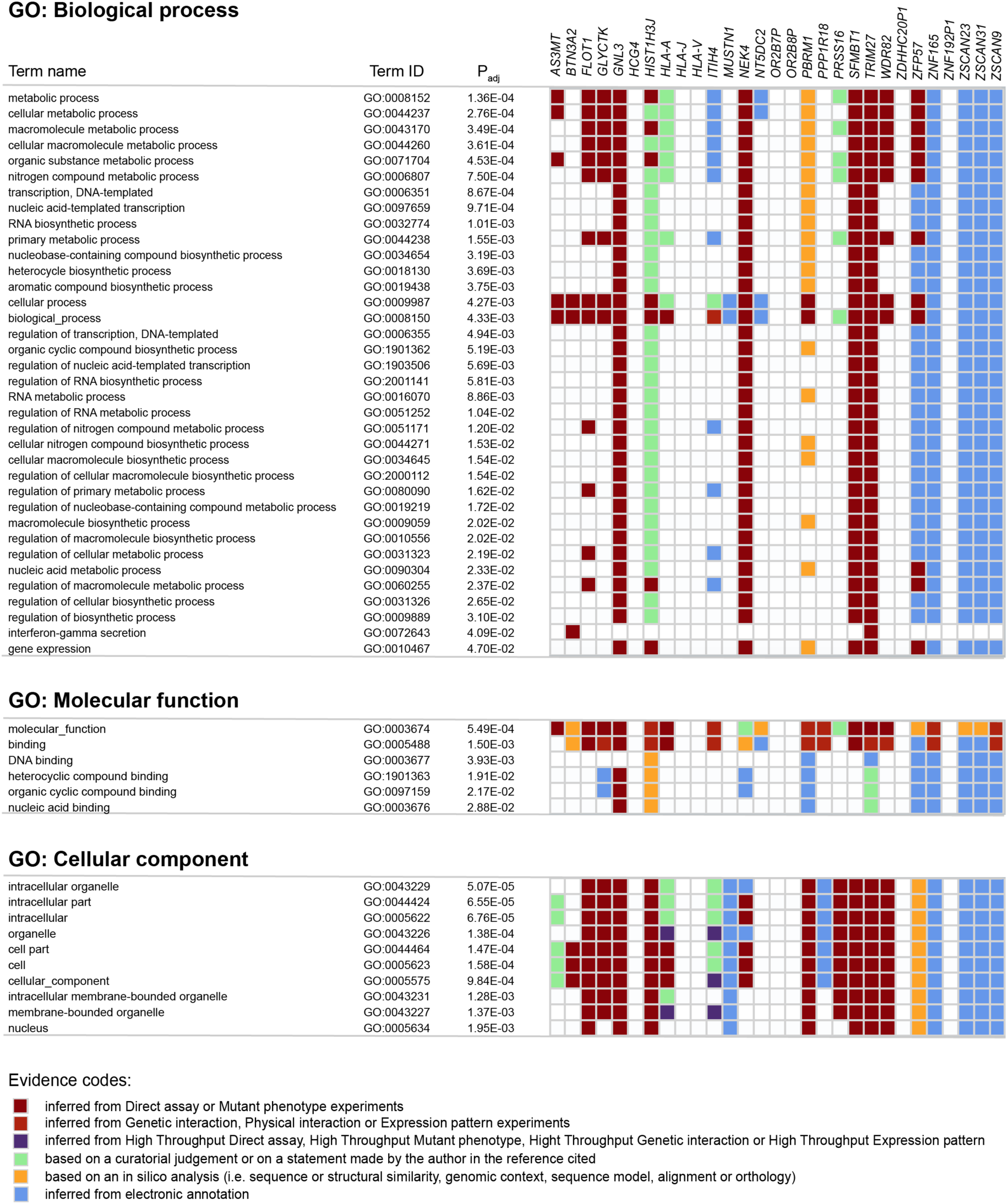
Functional enrichment analysis for 29 shared eGenes (4 eGenes are not present in the g:GOSt database) showed that most of them are enriched in “binding” with DNA and other heterocyclic and organic cyclic compounds. Protein products of the shared eGenes are also involved in gene expression, transcription, interferon-gamma secretion and various metabolic, biosynthetic and regulatory processes within a cell. The adjusted *p* < 0.05 determines the significance level for GO terms.

**Supplementary Fig. 9.**
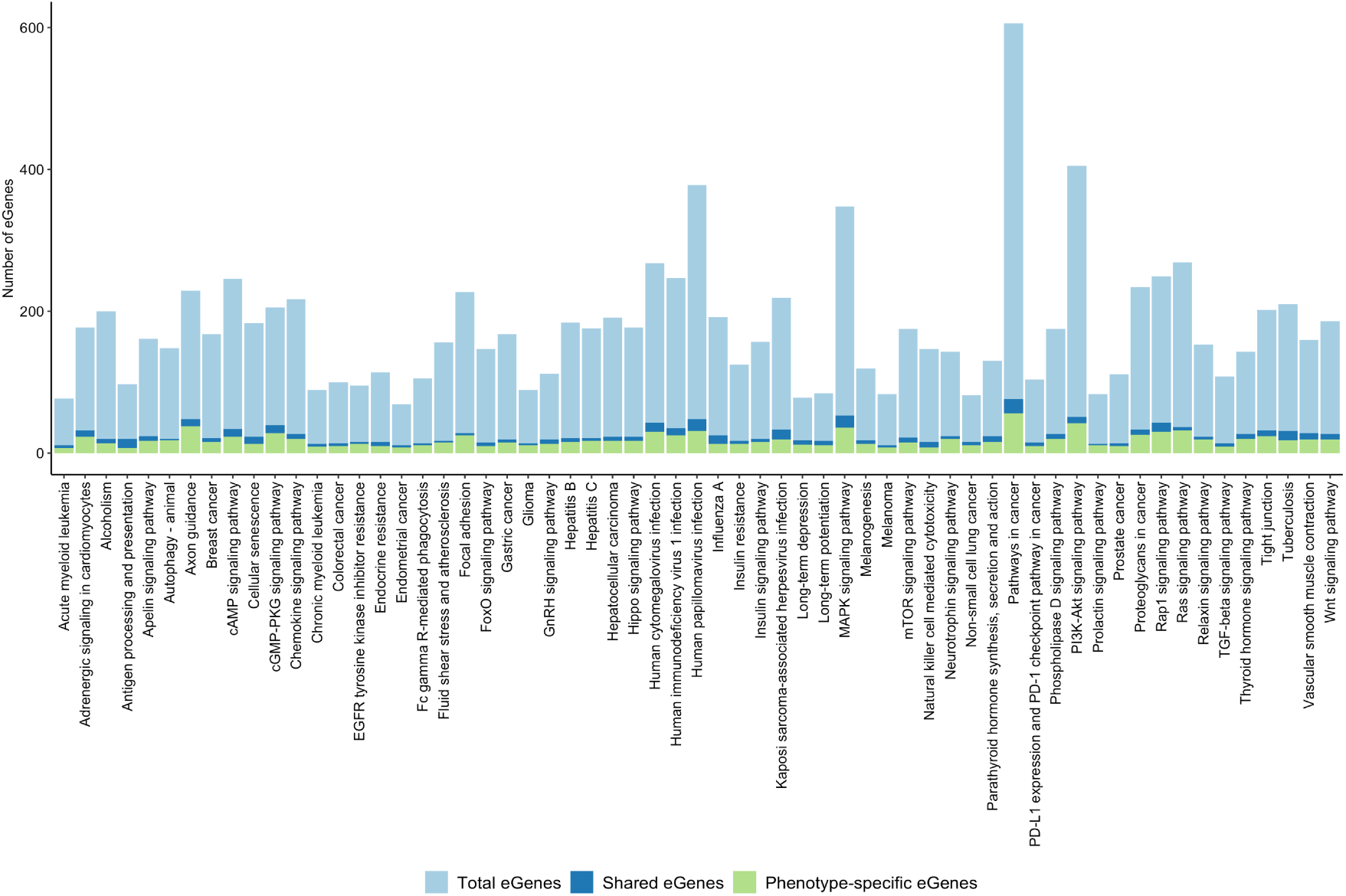
Phenotype-specific eGenes and eGenes shared by at least two phenotypes are co-occurring within 61 shared biological pathways.

**Supplementary Fig. 10.**
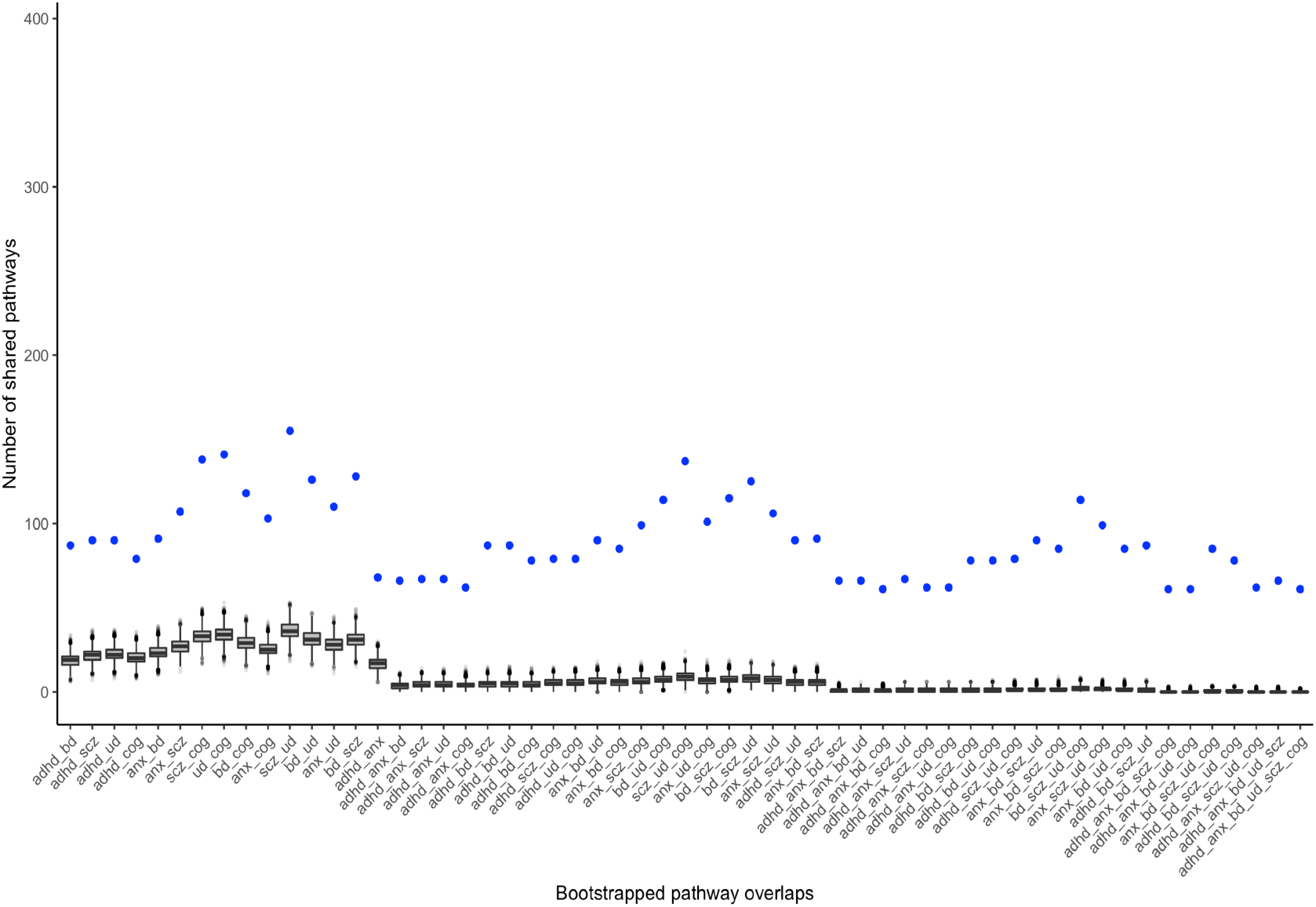
Box plots of the bootstrapped (10,000 iterations) pathway overlaps among psychiatric disorders and cognition. Blue dots indicate the number of shared pathways in the observed overlaps. The bootstrap test shows that the pathways overlaps are statistically significant (*p* < 0.01) and didn’t occur due to chance.

**Supplementary Fig. 11.**
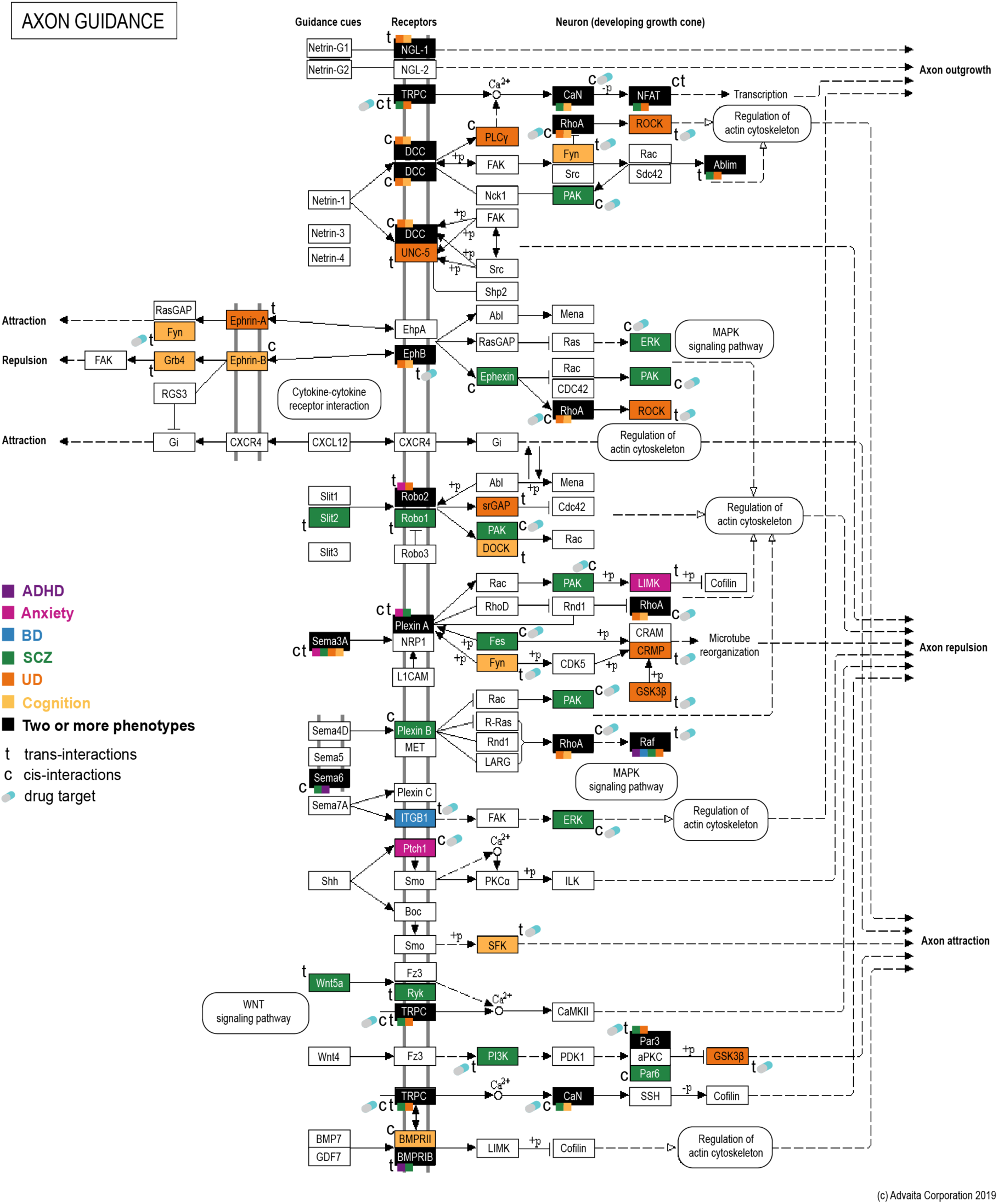
SNPs mark eQTLs that act to regulate genes within the axon guidance pathway. Axon guidance represents a key stage in the development of neuronal network. Axons are guided by netrins, ephrins, Slits, semaphorins and other guidance factors to reach their correct targets and form precise functional circuits. Co-occurrence of shared and phenotype-specific affected eGenes in this pathway may lead to a series of cellular events associated with dysregulation and disintegration of these circuits in psychiatric and cognitive phenotypes.

**Supplementary Fig. 12.**
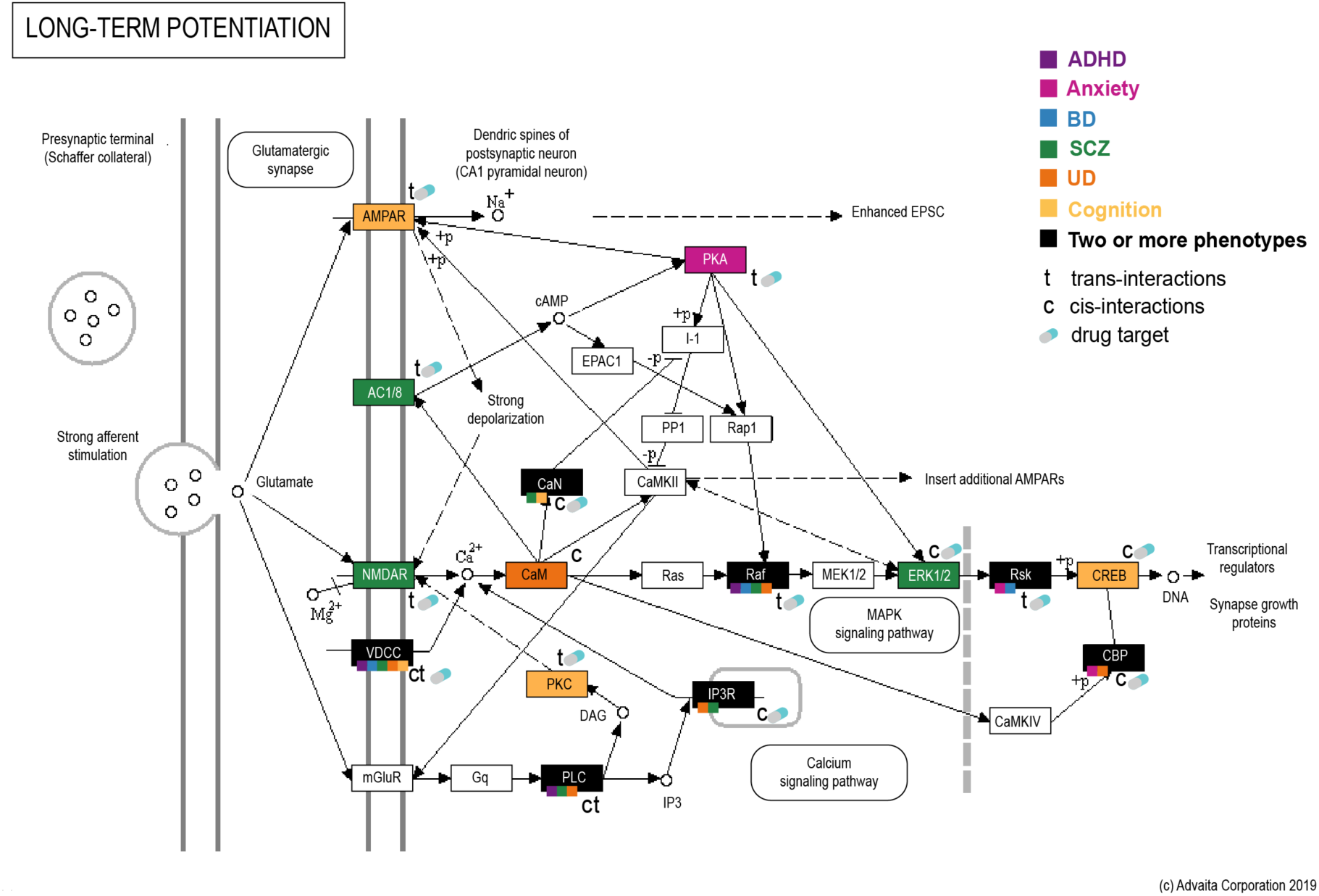
Co-occurrence of genes regulated by phenotype-associated eQTLs within the long-term potentiation (LTP) pathway - the molecular basis for learning and memory. Impaired LTP may have a role in psychiatric disorders and cognitive functioning.

**Supplementary Fig. 13.**
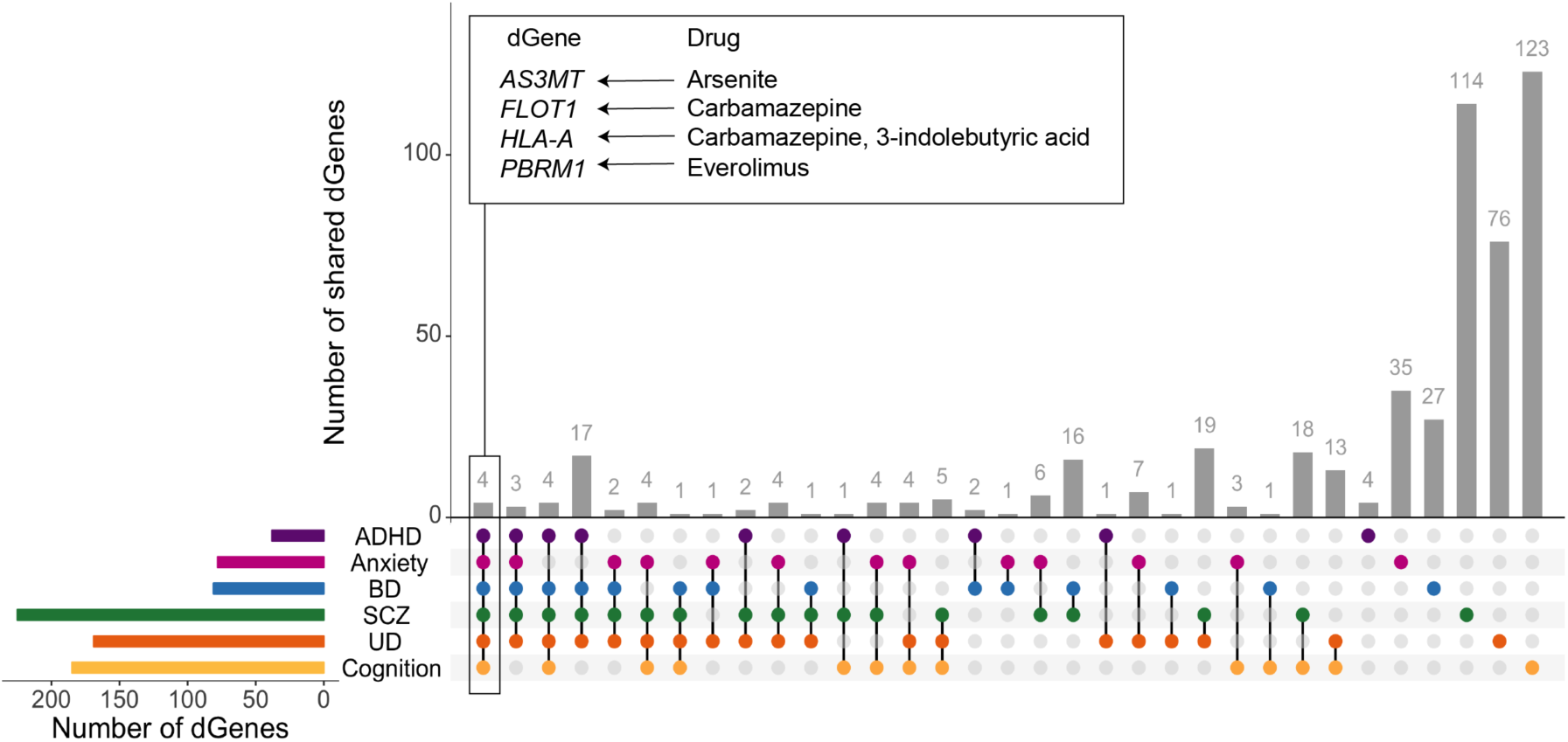
Psychiatric disorders and cognition share draggable eGenes (dGenes) whose products represent potential drug targets. Four dGenes (*AS3MT*, *FLOT1*, *HLA-A* and *PBRM1*) are shared across all six phenotypes.

**Supplementary Fig. 14.**
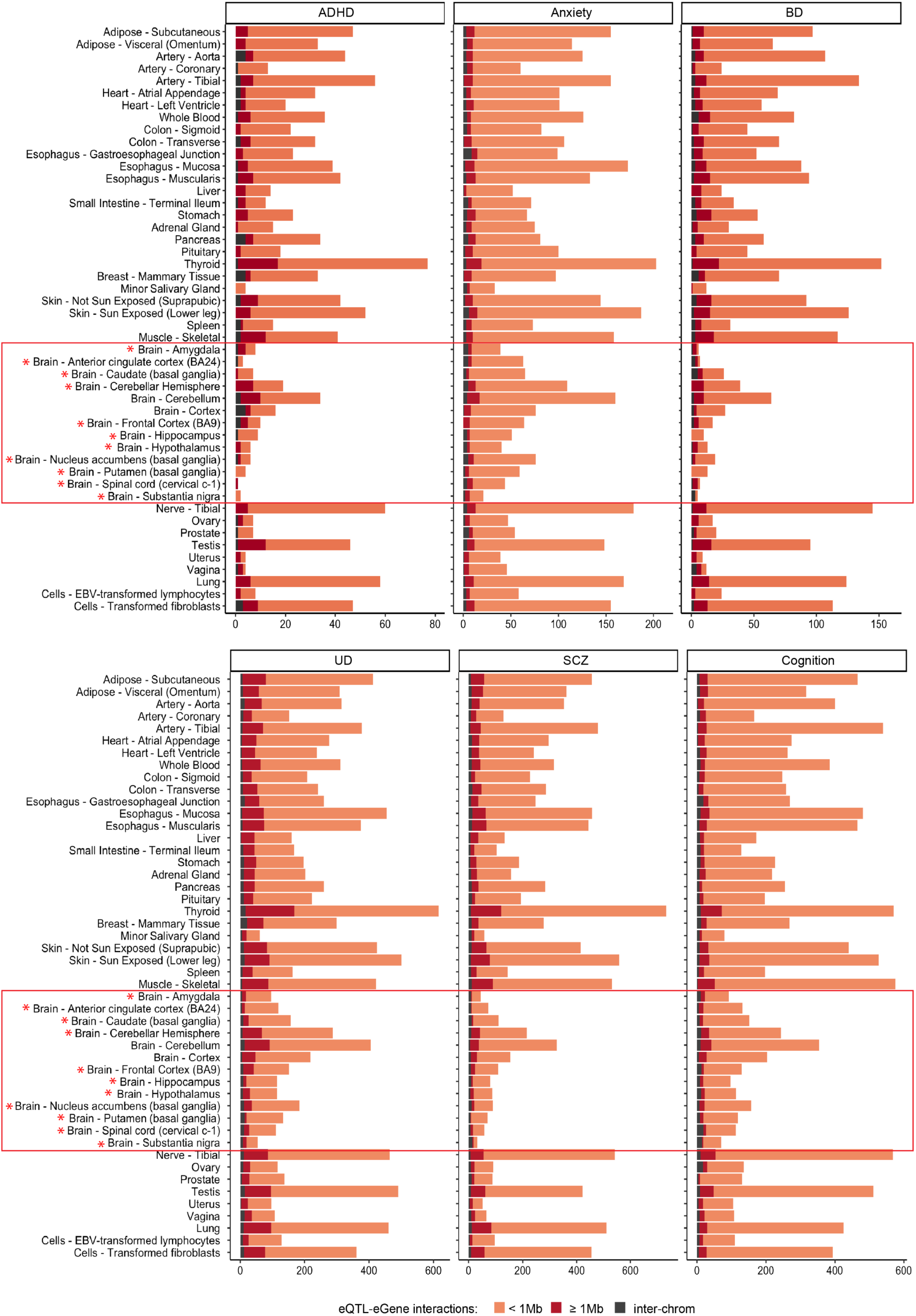
Number of tissue-specific eQTL SNP-eGene interactions identified for psychiatric disorders and cognition. Red asterisks specify brain tissues that were re-sampled and have a longer ischemic time (the time from death until the time of the sample fixation) compared to other tissues.

**Supplementary Fig. 15.**
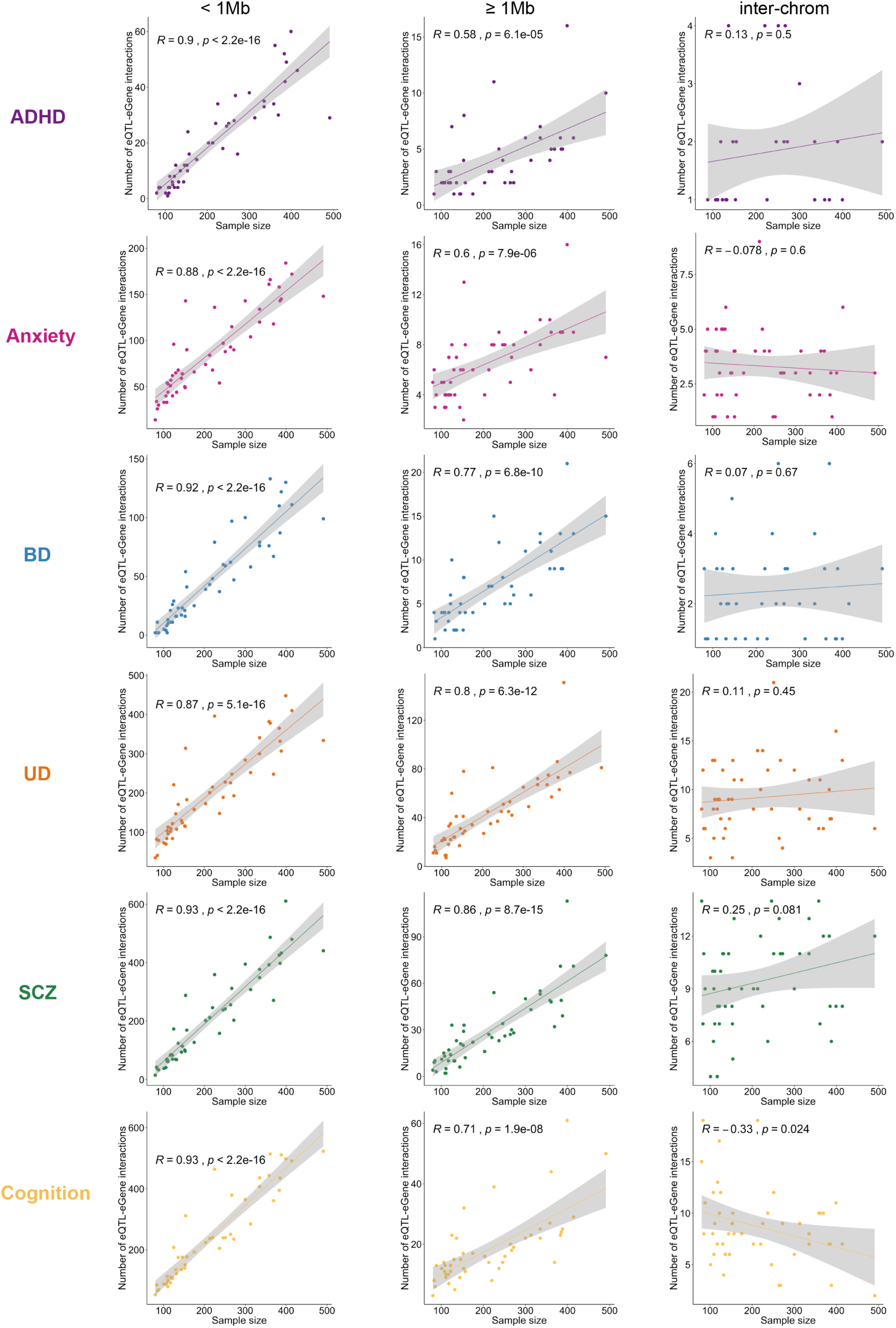
Correlation analysis between GTEx tissue sample size and the number of cis- and trans-acting eQTL SNP-eGenes interactions identified for psychiatric disorders and cognition. Shaded regions represent 95% confidence intervals containing the true correlation. The larger is GTEx sample size, the more <1Mb and ≥1Mb eQTL SNP-eGenes interactions are observed. However, the number of interchromosomal interactions tends to be not dependent on the GTEx sample size. For more information on cis-(<1Mb) and trans-acting (≥1Mb and interchromosomal) eQTL SNP-eGene interactions per GTEx tissue, see Supplementary Table 18.

